# The virome of the invasive Asian bush mosquito *Aedes japonicus* in Europe

**DOI:** 10.1101/2022.11.26.518030

**Authors:** Sandra R. Abbo, João P. P. de Almeida, Roenick P. Olmo, Carlijn Balvers, Jet S. Griep, Charlotte Linthout, Constantianus J. M. Koenraadt, Bruno M. Silva, Jelke J. Fros, Eric R. G. R. Aguiar, Eric Marois, Gorben P. Pijlman, João T. Marques

**Author notes:** Corresponding author. Corresponding authors |. Contributed equally (Sandra R. Abbo, João P. P. de Almeida).

## Abstract

**Background:** The Asian bush mosquito *Aedes japonicus* is rapidly invading North America and Europe. Due to its potential to transmit multiple pathogenic arthropod-borne (arbo)viruses including Zika virus, West Nile virus and chikungunya virus, it is important to understand the biology of this vector mosquito in more detail. In addition to arboviruses, mosquitoes can also carry insect-specific viruses that receive increasing attention due to their potential effects on host physiology and arbovirus transmission. In this study, we characterized the collection of viruses, referred to as the virome, circulating in *Ae. japonicus* populations in the Netherlands and France.

**Results:** Applying a small RNA-based metagenomic approach to *Ae. japonicus*, we uncovered a distinct group of viruses present in samples from both the Netherlands and France. These included one known virus, *Ae. japonicus* narnavirus 1 (AejapNV1), and three new virus species that we named *Ae. japonicus* totivirus 1 (AejapTV1), *Ae. japonicus* anphevirus 1 (AejapAV1) and *Ae. japonicus* bunyavirus 1 (AejapBV1). We also discovered sequences that were presumably derived from two additional novel viruses: *Ae. japonicus* bunyavirus 2 (AejapBV2) and *Ae. japonicus* rhabdovirus 1 (AejapRV1). All six viruses induced strong RNA interference responses, including the production of 21 nucleotide sized small interfering RNAs, a signature of active replication in the host. Notably, AejapBV1 and AejapBV2 belong to different viral families, however, no RNA-dependent RNA polymerase sequence has been found for AejapBV2. Intriguingly, our small RNA-based approach identified a ∼1 kb long ambigrammatic RNA that is associated with AejapNV1 as a secondary segment but showed no similarity to any sequence in public databases. We confirmed the presence of AejapNV1 primary and secondary segments, AejapTV1, AejapAV1 and AejapBV1 by reverse-transcriptase PCR in wild-caught *Ae. japonicus* mosquitoes. AejapNV1 and AejapTV1 were found at high prevalence (87-100%) in adult females, adult males and larvae.

**Conclusions:** Using a small RNA-based, sequence-independent metagenomic strategy, we uncovered a conserved and prevalent virome among *Ae. japonicus* mosquito populations. The high prevalence of AejapNV1 and AejapTV1 across all tested mosquito life stages suggests that these viruses are intimately associated with *Ae. japonicus* and may affect different aspects of the physiology of this vector mosquito.

## BACKGROUND

Mosquitoes of the *Aedes* genus are responsible for mosquito-borne viral disease outbreaks worldwide. The tropical yellow fever mosquito *Aedes aegypti* and the invasive Asian tiger mosquito *Aedes albopictus* pose a large threat to human health by transmitting medically important arthropod-borne (arbo)viruses including Zika virus (ZIKV), chikungunya virus (CHIKV), dengue virus (DENV) and yellow fever virus [1, 2]. Research has specially focused on these two urban mosquito species, however, also other *Aedes* species are becoming increasingly widespread [3–5] and their role in arbovirus transmission requires further attention.

The Asian bush mosquito *Aedes japonicus*, which originates in Northeast Asia, is a highly invasive species [6]. During the past two decades, this mosquito species quickly spread to North America and Europe, where it established large, permanent populations despite intensive control efforts [6, 7]. Importantly, *Ae. japonicus* is capable of transmitting multiple arboviruses including West Nile virus [8, 9], Japanese encephalitis virus [10], ZIKV [11, 12], Usutu virus (USUV) [12] and CHIKV [13]. Therefore, it is important to understand the biology of this exotic mosquito species in greater detail.

In addition to arboviruses, mosquitoes can also carry insect-specific viruses (ISVs). Whereas arboviruses are maintained in transmission cycles between arthropods and vertebrate animals or humans, ISVs do not replicate in vertebrates, and only infect insects. ISVs can persistently infect mosquito populations and have recently received increasing attention due to their potential effects on host physiology and arbovirus transmission [14–18]. Exploring the diversity of ISVs may also help to better understand the evolution of arboviruses and to develop new strategies for arbovirus control [19].

Extensive virome analyses that have been performed for *Ae. aegypti* and *Ae. albopictus* [18, 20–22] showed that both mosquito species harbour a large diversity of ISVs. For *Ae. japonicus*, however, an in-depth virome analysis is still missing, although a first glimpse has revealed the presence of a novel narnavirus in this mosquito. This *Ae. japonicus* narnavirus 1 (AejapNV1) was discovered in *Ae. japonicus* from the Netherlands [12] and was also found in *Ae. japonicus* from Japan shortly thereafter [23], indicating its close association with *Ae. japonicus*. AejapNV1 belongs to a novel group of unique, ambigrammatic narnaviruses, which not only contain an open reading frame (ORF) coding for the RNA-dependent RNA polymerase (RdRp) on the positive strand but also a very long ORF with unknown function on the (reverse-complement) negative strand [24, 25].

Metagenomic approaches have played an essential role in uncovering the virome of vector mosquitoes and the discovery of many novel ISVs [15]. Despite the success of these approaches, certain aspects of virome analysis remain challenging to fulfil using large-scale nucleic acid sequencing, such as: the detection and classification of highly divergent viral sequences that do not align to any known reference sequence (i.e., the viral ‘dark matter’), the association of sequences from different genomic segments of the same virus, and the differentiation of exogenous viruses from endogenous viral elements (EVEs).

These limitations can be overcome using small RNA-based metagenomics. Virus-derived small RNAs are produced in mosquitoes when viral double-stranded RNA (dsRNA) replication intermediates are recognized and cleaved by Dicer-2, a key protein of the antiviral RNA interference (RNAi) machinery, which results in the production of 21 nucleotide (nt) small interfering RNAs (siRNAs) [26]. The formation of viral siRNAs is a signature of active virus replication [27]. In addition to viral siRNAs, viral PIWI-interacting RNAs (piRNAs) of ∼24-29 nt in length are also produced for some viruses [26]. Thus, small RNA sequencing detects products derived from host antiviral pathways that discriminates EVEs and allows for sequence-independent characterization of viral sequences [28].

In the current study, we applied a small RNA-based metagenomic approach to analyze the virome of wild-caught *Ae. japonicus* mosquitoes from populations in the Netherlands and France, and identified AejapNV1 and three novel virus species. Interestingly, for AejapNV1, not only the primary genome segment (S1) encoding the RdRp was found, but we also discovered an ambigrammatic secondary genome segment (S2) with unknown function. S2 did not align to any known sequence in public databases. Still, it could be associated with the same virus on the basis of similar (di)nucleotide composition in S1 and S2, and by applying unsupervised clustering techniques using small RNA features generated by our small RNA metagenomic approach. Additionally, we found sequences derived from two putative, novel virus species, and together with the other four discovered viruses, we characterized their genome organisation and small RNA profiles. Finally, reverse transcriptase PCR (RT-PCR) was used to study the prevalence of these viruses in *Ae. japonicus* adults, larvae and eggs.

## METHODS

### Small RNA library preparation and high-throughput sequencing

*Ae. japonicus* adult female mosquitoes from Strasbourg, France, were collected using human landing catches (HLCs) [12] and further identified using morphological characteristics. Two pools were made prior to RNA extraction, one containing two (FR_01) individuals and another containing four (FR_02) as shown in **Fig. 1A**. Total RNA was isolated using TRIzol (Invitrogen) following the manufacturer’s protocol. Briefly, mosquitoes were transferred to 1.5 ml screw-capped tubes containing ceramic beads (1.4 mm in diameter, Omini) and ice-cold TRIzol (Invitrogen). Mosquitoes were grinded using a Precellys Evolution Homogenizer at 6,500 rpm in 3 cycles of 20 seconds each. Then, 10 µg of glycogen (Ambion) was added to the aqueous layer of each sample to facilitate pellet visualization upon RNA precipitation. RNAs were resuspended in RNAse-free water (Ambion) and stored at −80°C until further notice. Total RNA was used as input for library preparation utilizing the kit NEBnext Multiplex Small RNA Library Prep Set for Illumina following the recommended protocol, except for one minor modification: the 5’ adapter was replaced by an analogue that contains 6 extra nucleotides at the 3’ extremity to improve barcoding precision, which is sequenced along with the cloned small RNA. Libraries were sequenced at the GenomEast sequencing platform at the Institut de Génétique et de Biologie Moléculaire et Cellulaire in Strasbourg, France, using Illumina HiSeq 4000 equipment. The *Ae. japonicus* small RNA libraries from Lelystad, the Netherlands, reanalyzed in this study, were previously deposited in the NCBI sequence read archive (SRA) under BioProject PRJNA545039 [12]. Libraries NL_01 and NL_02 were built from pools of four and six adult females, respectively (**Fig. 1A**).

**Figure 1.**
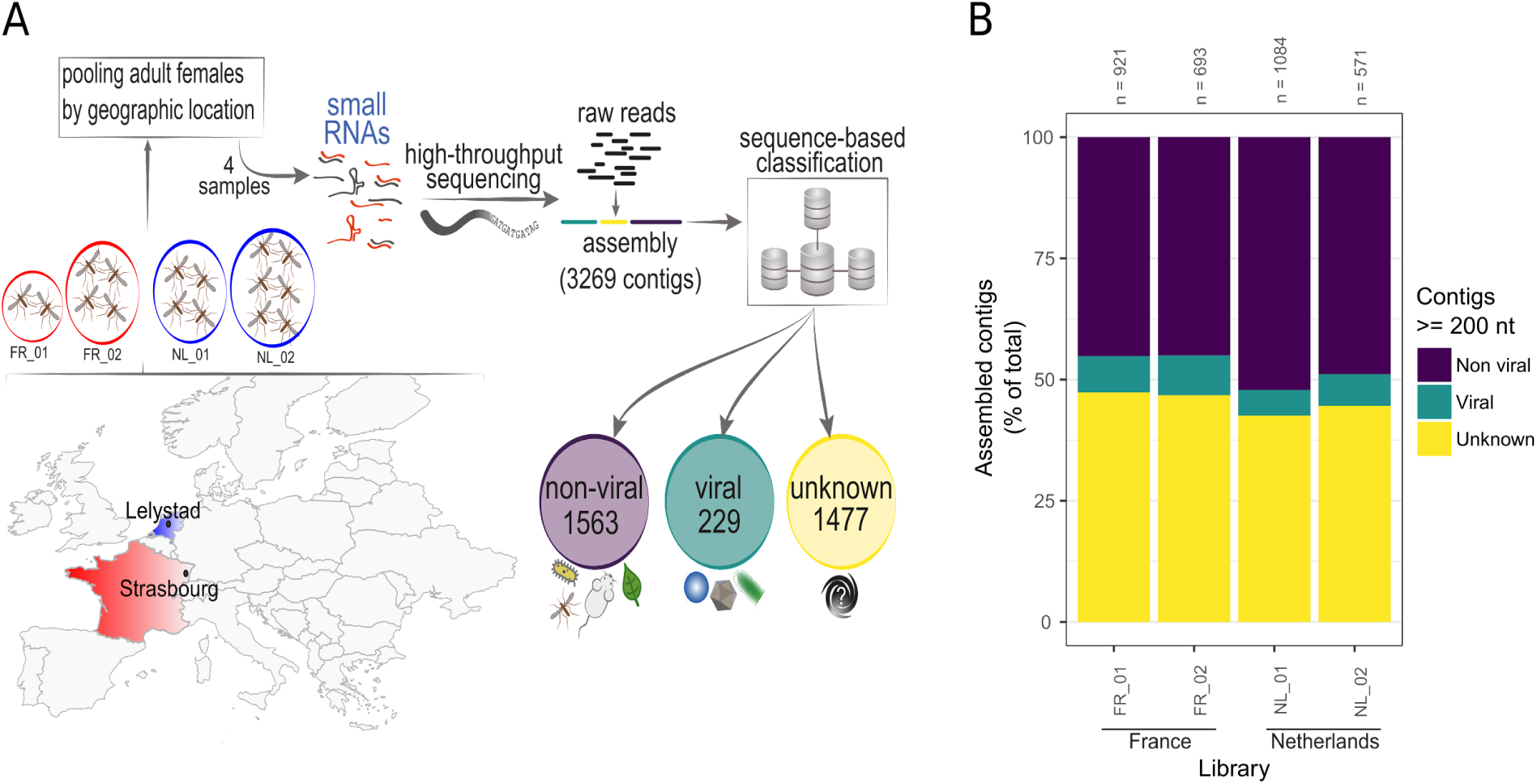
Analysis of the *Ae. japonicus* virome using a small RNA-based metagenomic approach. **(A)** Map of Europe indicating mosquito collection sites: Strasbourg, France (red) and Lelystad, the Netherlands (blue). Pools with the number of sampled mosquitoes from Strasbourg are indicated inside the red circles (FR_01 two mosquitoes and FR_02 four mosquitoes) and from Lelystad inside the blue circles (NL_01 four mosquitoes and NL_02 six mosquitoes). Captured mosquitoes were morphologically identified by species. Samples were used to prepare small RNA libraries for high-throughput sequencing. Sequencing results were analyzed using our metagenomic pipeline. Assembled contigs were classified into non-viral, viral, and unknown sequences based on sequence similarity against reference databases. **(B)** Individual results from our sequence similarity analysis for each of the four small RNA libraries in this study. The total number of contigs bigger than or equal to 200 nt (n) and the proportion of non-viral, viral and unknown contigs are shown.

### Small RNA-based metagenomics for virus identification

Raw reads from small RNA libraries were submitted to adapter trimming using Cutadapt v1.12 [29], and Illumina libraries from France had the 6 inserted nucleotides trimmed with the adapters. Sequences with an average Phred quality below 20, ambiguous nucleotides, and/or a length shorter than 15 nt were discarded. The remaining sequences were mapped to genome sequences of *Ae. aegypti* (AaeL5) [30], *Ae. albopictus* [31], and bacterial reference genomes using Bowtie v1.3 [32]. Unmapped reads from the previous step were assembled in contigs combining metaSPAdes [33] and Velvet v1.0.13 [34]. More details about k-mers and read sizes combinations for assemblies using small RNAs in the current work can be found in [28]. Contigs larger than 200 nt were characterized based on sequence similarity against the NCBI *nt* database using BLAST+ [35] and against *nr* using DIAMOND (mode *--very-sensitive*) [36], considering significant hits with *e-values* lower than 1e^-5^ for nucleotide comparison or 1e^-3^ for amino acid comparison. For small RNA size profile and coverage analysis, reads unmapped to mosquito and bacterial genomes were aligned to viral and unknown contigs using Bowtie v1.3, allowing one mismatch. Size profiles of small RNAs matching reference sequences and 5’ nt base preference were calculated using in-house Perl v5.16.3, BioPerl library v1.6.924 and R v4.0.5 scripts (https://github.com/JPbio/small_RNA_MetaVir/). Plots were generated in R using ggplot2 v3.3.5 package. For manual curation of putative viral contigs, top five BLAST and DIAMOND hits were analyzed to rule out the similarity to other organisms; ORF organization and small RNA profiles (size distribution and coverage) were analyzed to differentiate exogenous from endogenous viruses and check for inconsistencies. Contigs containing truncated ORFs and small RNA profiles without symmetric small RNA peaks at 21 nt were considered putative EVEs as described [26]. For more details on manual curation steps, see [18]. Redundancy of curated viral contigs was removed using CD-HIT [37] requiring 90% coverage of the smaller sequence (*-aS*) with 90% of global identity (*-c*). Representative unique contigs from the previous step were used for co-occurrence analysis based on small RNA abundancy in each of the small RNA libraries using Reads Per Kilobase of transcript per Million reads (RPKM) values for reads ranging from 20 to 22 nt aligned to the viral and unknown contigs. Hierarchical contig clusters using RPKM values were constructed in R using Euclidean distance and average method. Z-scores were calculated based on the frequency of each small RNA size from 15 to 35 nt mapped to representative viral and unknown contigs considering each strand polarity separately using the library where the contig was originally assembled. Strand polarity for contig comparisons was adjusted based on the reads abundance per strand and coding ORFs direction of each contig. Heatmaps for RPKM and Z-score values were plotted using the package ComplexHeatmap in R [38]. As an attempt of viral sequence extension as proposed by Sardi et al., 2020 [39], contigs grouped in the same cluster with sequence similarity hits to the same virus were submitted to a new assembly round using SPAdes [40] with the parameter *--trusted-contigs* using all the libraries in which that viral sequences were found.

### piRNA analysis

Sequence overlaps and signatures of viral-derived small RNAs were analysed to determine the presence of piRNAs. Distances between small RNAs were calculated as previously described [41, 42]. Briefly, the frequency of 5′ to 5′ distances between 24 to 29 nt long reads from opposite strands was calculated and normalized by Z-score. A-U enrichment for 1st and 10th positions of 24 to 29 nt long reads was observed by filtering these reads after alignment to viral sequences and plotting sequence logos with the ggseqlogo R package [43].

### Phylogenetic analysis

We selected the contig containing the largest RdRp sequence for the identified viruses. For bunyaviruses, we also selected the contigs containing the largest sequences of nucleocapsid and glycoprotein. The presence of conserved protein domains for RdRp, nucleocapsid, and glycoprotein was confirmed with NCBI Conserved Domain Search (https://www.ncbi.nlm.nih.gov/Structure/cdd/wrpsb.cgi). Searches for potential homologous viral sequences were performed using Blastx against the *nr* database. Coding sequences were translated to amino acids, and multiple sequence alignments were performed using MAFFT [44]. The best-fit protein evolution model was selected using MEGA-X [45] under the Akaike Information Criterion. Phylogenetic inference was executed with MEGA-X using Maximum Likelihood method. For all phylogenetic trees, clade robustness was assessed using the bootstrap method (1000 pseudoreplicates). The trees were viewed using iTOL version 6.5 [46].

### Analyses of CpG dinucleotide frequency and GC content

Mononucleotide frequencies, dinucleotide frequencies and the observed/expected (O/E) ratios of viral and unknown sequences were determined using the function Composition Scan within SSE version 1.4 [47].

### Sequence alignment and RNA structure modelling

Narnavirus RNA sequences were aligned using MUSCLE version 3.8.1551 [48]. Narnavirus RNA secondary structures were predicted using the RNAstructure web server version 6.3 with temperature set to 28°C [49]. Pseudoknots in totivirus RNA were predicted with DotKnot version 1.3.2 [50]. RNA folding was visualised using VARNA [51].

### Protein structure analysis

Primary structure physical-chemical properties were calculated with ProtParam software [52]. Global sequence alignments were performed with Clustal Omega [53] and colored with ESPcript web server [54]. PSIPRED Workbench [55] was used for structural assessments, with secondary structures predicted using PSIPRED 4.0 [56], DISOPRED3 [57] for disordered regions and MEMSAT-SVM [58] for membrane interaction. AlphaFold [59] tertiary structure modelling was performed locally with the complete database downloaded and using default parameters and “06/01/2022” as maximum template date.

### Mosquito collection and rearing for RT-PCR

To obtain mosquito samples for RT-PCR analyses, *Ae. japonicus* mosquitoes were collected in Lelystad, the Netherlands using oviposition traps, water reservoirs in local rain barrels and HLCs as described [12] during the summer of 2020. Adult females obtained from HLCs were stored at −80°C until further analysis, whereas eggs and larvae were kept in the laboratory at 23°C and 60% relative humidity. Adult males were grown from collected eggs and larvae as described [12], and subsequently stored at −80°C. Pools of 25 eggs or individual fourth instar larvae were also stored at −80°C. The common house mosquito *Culex pipiens* (biotype *pipiens*) was used as negative control for the RT-PCR analyses. *Cx. pipiens pipiens* mosquitoes collected in Wageningen, the Netherlands during the summer of 2020, were used to establish a laboratory colony as previously described [60]. Mosquitoes were fed with chicken whole blood (Kemperkip, Uden, the Netherlands) through Parafilm using a Hemotek PS5 feeder (Discovery Workshops, Lancashire, United Kingdom), and reared at 23°C and 60% relative humidity. Individual adult female mosquitoes were stored at −80°C for further analysis.

### Mosquito RNA isolation for RT-PCR

Frozen mosquitoes were homogenized using a Bullet Blender Storm (Next Advance, Averill Park, NY, USA) in combination with 0.5 mm zirconium oxide beads (Next Advance; for adult mosquitoes and larvae) or 0.9-2.0 mm stainless steel beads (Next Advance; for eggs) at maximum speed for 2 min. Afterwards, homogenates were centrifuged at maximum speed in an Eppendorf 5424 centrifuge for 1 min. Pellets were resuspended in 450 μl lysis buffer from the innuPREP DNA/RNA Mini Kit (Analytik Jena, Jena, Germany) and subsequently incubated for 20 min to lyse. RNA was isolated using the innuPREP DNA/RNA Mini Kit (Analytik Jena) according to manufacturer’s instructions. RNA yields were measured using a NanoDrop ND-1000 spectrophotometer.

### RT-PCR analysis

Viruses were detected by RT-PCR using a 2720 Thermal Cycler (Applied Biosystems, Foster City, CA, USA) and SuperScript III One-Step RT-PCR System with Platinum *Taq* DNA polymerase (Invitrogen) according to manufacturer’s protocol. In the case of a dsRNA virus, total RNA was subjected to a 5 min incubation step at 95 °C prior to the RT-PCR reaction to denature the viral dsRNA. 50 ng of mosquito total RNA was used as input for RT-PCR, and each virus was amplified using specific primers targeting the viral RdRp sequence (**Table 1**). For AejapNV1, primers were not only designed for the primary segment (RdRp), but also for the secondary segment (**Table 1**). To test the RNA quality of the samples, RT-PCRs against mosquito ribosomal protein S7 RNA were also included (**Table 1**). The RT-PCR product of the secondary segment of AejapNV1 was sent for DNA sequencing to Macrogen (EZ-Seq protocol).

**Table 1.**
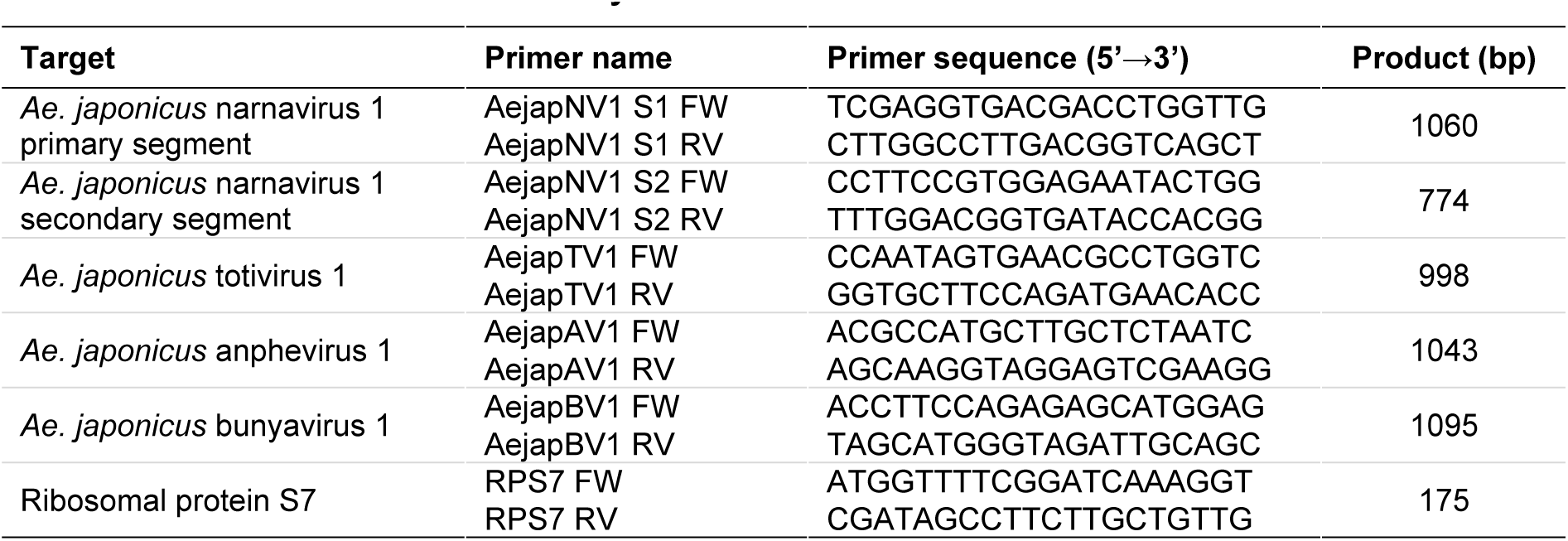
Primer sets used in this study.

## RESULTS

### Classification of contigs based on sequence similarity

To assess the collection of viruses found in *Ae. japonicus*, two new small RNA libraries of *Ae. japonicus* from France (FR_01 and FR_02) were sequenced and two published small RNA libraries of *Ae. japonicus* from the Netherlands (NL_01 and NL_02) [12] were reanalyzed (**Fig. 1A**; **Suppl. Table 1**). Libraries were processed and analyzed following a small RNA-based metagenomic strategy optimized to detect viral sequences [18]. General information on small RNA libraries and SRA accessions can be found in **Suppl. Table 1**.

In total, 3,269 contiguous sequences (contigs) larger than 200 nt were assembled from the four individual libraries. A summary of the assembly results is shown in **Suppl. Table 2**. Based solely on sequence similarity searches against NCBI non-redundant nucleotide and protein databases (*nt* and *nr*), contigs were classified into 229 viral, 1,563 non-viral, and 1,477 unknown sequences (**Fig. 1A**; **Suppl. Table 3**). Proportions of each class per library are shown in **Fig. 1B**.

### Curation of viral contigs

EVEs are often misperceived as viruses in metagenomic studies based on RNA sequencing (also referred to as metatranscriptomics) and this can overestimate the number of viruses in a given sample. To discriminate sequences derived from viruses and EVEs, we used a previously established analysis pipeline [18]. Briefly, this pipeline relies on investigating the small RNA profile and ORF composition of each contig previously classified as of viral origin by sequence similarity. Applying this filter, 93 sequences derived from exogenous viruses were obtained (**Suppl. Table 3**) out of the initial 229 viral contigs (**Suppl. Table 4**). The remaining 93 contigs were clustered by sequence similarity using CD-HIT to remove redundancy, resulting in 56 unique representative viral contigs (**Suppl. Table 5**). To evaluate only the natural circulating virome, 3 USUV and 14 ZIKV contigs from experimental infections assembled in the libraries NL_01 and NL_02 were removed, leaving a total of 39 unique representative sequences.

In an effort to associate different segments that may belong to the same virus, we evaluated co-occurrence of the viral contigs in the four libraries. In this regard, we determined the normalized number of small RNA reads that could be mapped to each of the

39 representative viral contigs. Here we also included 22 non-redundant contigs of unknown origin, that have lengths larger than 500 nt and passed the filters based on small RNA and ORF composition. Co-occurring contigs that shared a similar small RNA size profile were considered probable fragments from the same virus (**Fig. 2**). This analysis identified five distinct clusters of co-occurring contigs (**Fig. 2**). Notably, most of the contigs in each of the five clusters showed significant similarity to the same reference virus (**Suppl. Table 4**).

**Figure 2.**
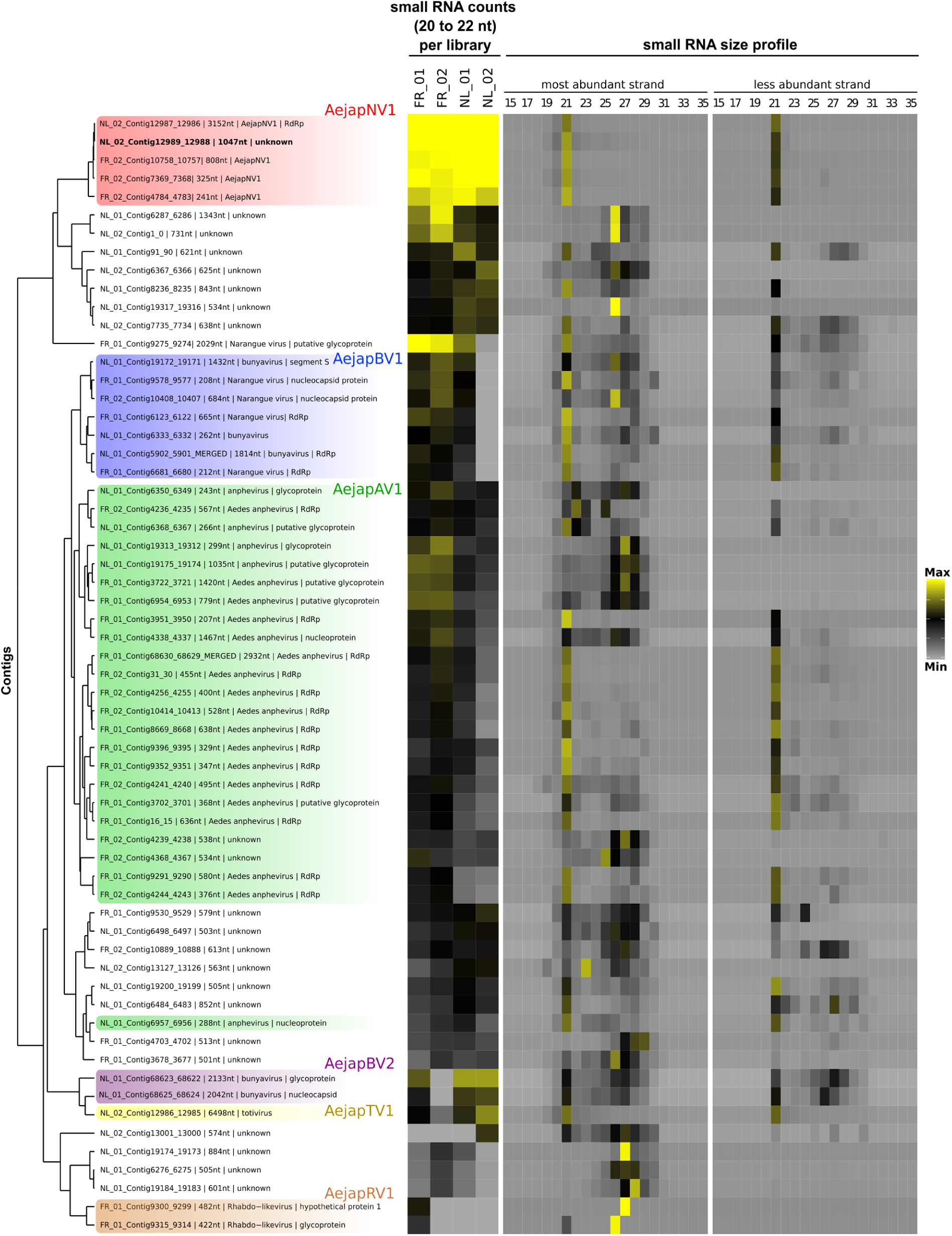
Co-occurrence of viral and unknown contigs. Hierarchical clustering of viral and unknown contigs assembled from small RNAs derived from *Ae. japonicus*. Clustering was based on Euclidean distance of RPKM values of small RNA counts with size from 20 to 22 nt in each library applying Average method. Contig clusters were defined using the dendrogram. Contigs inferred to be from the same virus were colored equally. Heatmap on the left represents the small RNA abundance for each curated contig in *Ae. japonicus* libraries. We plotted the Log2 of the RPKM values of small RNA counts with sizes from 20 to 22 nt (maximum value: 10; minimum value: 0). Heatmap on the right represents Z-score values for small RNAs from each size from 15 to 35 nt divided by strand polarity in the library in which the contig was originally assembled (maximum value: 7; minimum value: −1).

The first cluster, highlighted in red, contained four contigs with high sequence similarity to AejapNV1 and an additional unknown contig (NL_02_Contig12989_12988) (**Fig. 2**). Highlighted in blue, there are contigs with significant similarity to different segments of Narangue virus, a bunyavirus (**Fig. 2**). These contigs form a consistent cluster, except for one contig with high similarity to a bunyavirus glycoprotein (FR_01_Contig9275_9274) that is out of this cluster. Another large cluster of sequences displayed significant similarity to *Aedes* anphevirus and included two unknown contigs (**Fig. 2**). Two contigs with significant sequence similarity to a bunyavirus (highlighted in purple) and another to a totivirus (highlighted in yellow) also formed one cluster (**Fig. 2**). Two contigs with high sequence similarity to a rhabdovirus also formed a clear cluster (highlighted in brown in **Fig. 2**). Other clusters were not highlighted because they were composed of unknown contigs that could not clearly be associated to a specific virus. Apart from ZIKV and USUV, which were introduced by artificial infection, no contigs showed similarity to known arboviruses.

### Virus identification

To further characterize putative viruses represented by the contigs from the clusters highlighted in **Fig. 2**, we focused on sequences encoding viral polymerases. It was possible to identify contigs encoding viral polymerases in four clusters that were compared to the most similar sequences in GenBank (**Table 2**). One cluster contained AejapNV1 (genus *Narnavirus*, family *Narnaviridae*) that had been previously identified as a possible ISV [12]. According to phylogenetic analysis (**Fig. 3**), this virus clustered with other mosquito-associated narnaviruses containing long reverse ORFs (rORFs) from the *Alphanarnavirus* clade [25].

**Figure 3.**
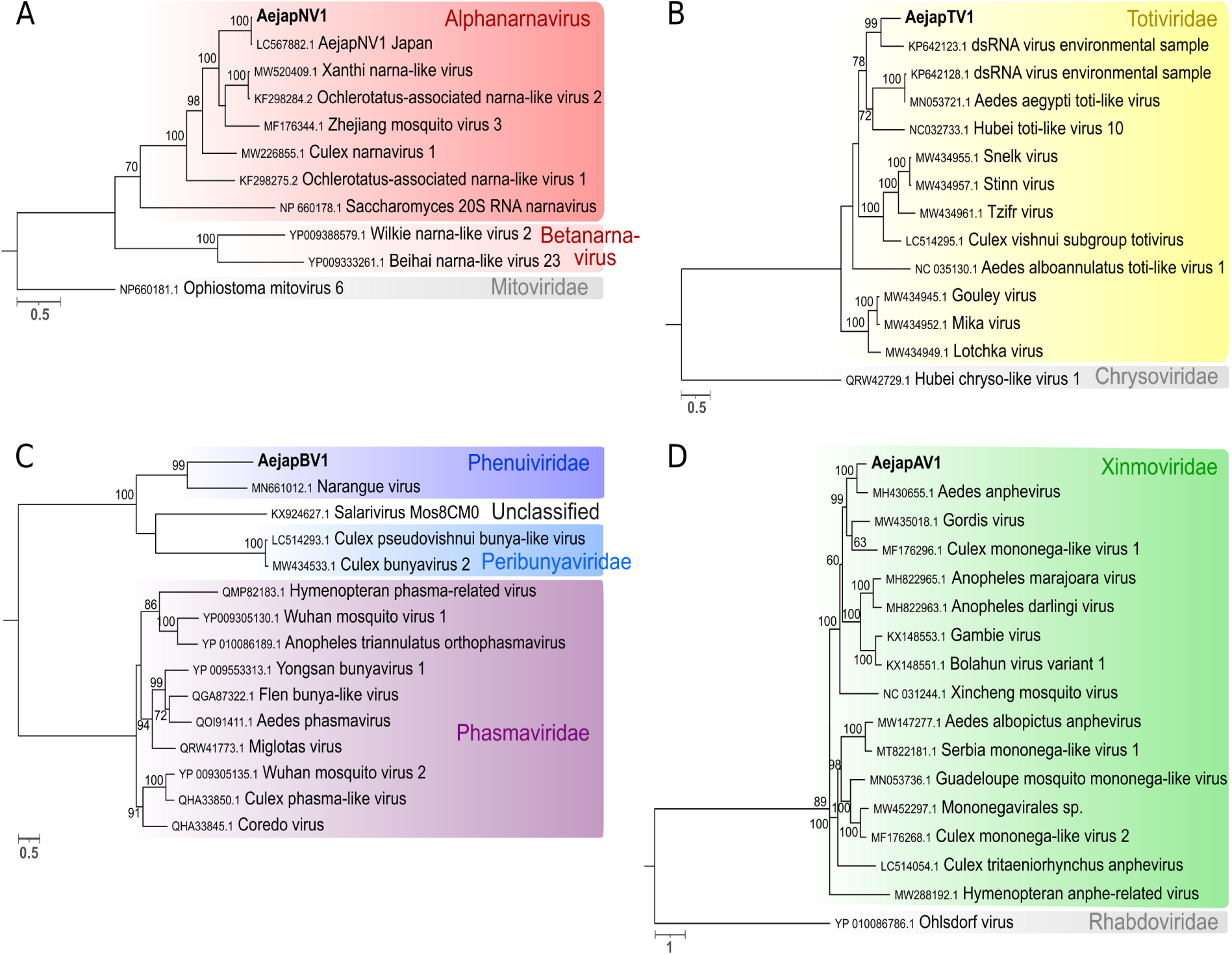
Phylogeny of viruses identified in *Ae. japonicus* mosquitoes. Phylogenetic trees were generated using the multiple sequence alignments of RdRp amino acid sequences. The trees were inferred by using the Maximum Likelihood method. The tree with the highest log likelihood is shown for each virus. Number of conserved sites and the substitution models used for each tree: **(A)** AejapNV1, 1446 sites, LG+G+F; **(B)** AejapTV1, 1269 sites, LG+G; **(C)** AejapBV1, 693 sites, LG+G+F; **(D)** AejapAV1, 1188 sites, LG+G+I+F. Node bootstraps were calculated with 1000 replicates and are shown close to each clade and values < 60 were omitted. Trees were midpoint-rooted, and RdRp sequences from distinct viral families were included in the alignments as outgroups. The trees are drawn to scale, branch lengths represent expected numbers of substitutions per amino acid site. Accession numbers for the nucleotide sequences from which the corresponding protein sequences were derived or the direct protein sequences are shown with the virus names. Viruses identified in this study are in bold.

**Table 2.**
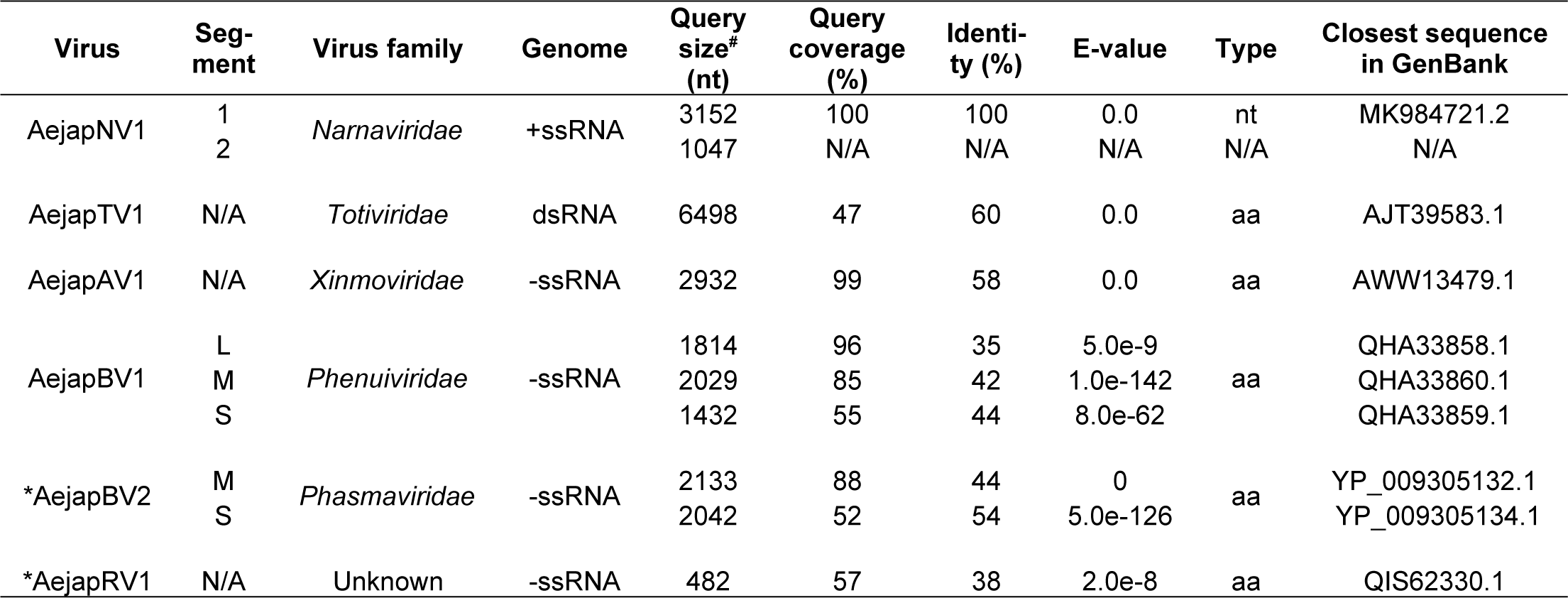
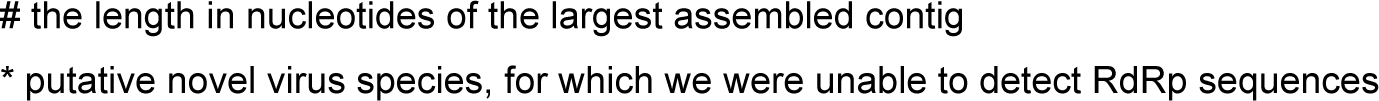
Viruses discovered by de novo assembly of small RNA reads from Ae. Japonicus mosquitoes collected in the Netherlands and France.

| Virus | Seg-<br>ment | Virus family | Genome | Query<br>size*<br>(nt) | Query<br>coverage<br>(%) | Identi-<br>ty (%) | E-value | Type | Closest sequence<br>in GenBank |
| --- | --- | --- | --- | --- | --- | --- | --- | --- | --- |
| AejapNV1 | 1 | *Narnaviridae* | +ssRNA | 3152 | 100 | 100 | 0.0 | nt | MK984721.2 |
|  | 2 |  |  | 1047 | N/A | N/A | N/A | N/A | N/A |
| AejapTV1 | N/A | *Totiviridae* | dsRNA | 6498 | 47 | 60 | 0.0 | aa | AJT39583.1 |
| AejapAV1 | N/A | *Xinmoviridae* | -ssRNA | 2932 | 99 | 58 | 0.0 | aa | AWW13479.1 |
| AejapBV1 | L | *Phenuiviridae* | -ssRNA | 1814 | 96 | 35 | 5.0e-9 | aa | QHA33858.1 |
|  | M |  |  | 2029 | 85 | 42 | 1.0e-142 |  | QHA33860.1 |
|  | S |  |  | 1432 | 55 | 44 | 8.0e-62 |  | QHA33859.1 |
| *AejapBV2 | M | *Phasmaviridae* | -ssRNA | 2133 | 88 | 44 | 0 | aa | YP_009305132.1 |
|  | S |  |  | 2042 | 52 | 54 | 5.0e-126 |  | YP_009305134.1 |
| *AejapRV1 | N/A | Unknown | -ssRNA | 482 | 57 | 38 | 2.0e-8 | aa | QIS62330.1 |

The other contigs containing viral polymerase sequences showed significant but low sequence similarity to known references of totiviruses, anpheviruses and bunyaviruses in GenBank only at the amino acid level (*e-value* < 1e^-3^) (**Table 2**). Phylogenetic analyses indicated that these are likely new viruses belonging to the *Totiviridae*, *Xinmoviridae*, and *Phenuiviridae* families that were named *Ae. japonicus* totivirus 1 (AejapTV1), *Ae. japonicus* anphevirus 1 (AejapAV1) and *Ae. japonicus* bunyavirus 1 (AejapBV1), respectively (**Fig. 3**). Based on comparisons to the closest reference, we observed that our assembly likely represented the full genome of AejapTV1 but that the genomes of AejapBV1 and AejapAV1 were not complete. In **Suppl. Fig. 1**, the putative genome organizations of AejapBV1 and AejapAV1 based on the reference sequences are shown. All three new viruses were closely related to known ISVs (**Fig. 3**; **Table 2**), but their final classification requires further biological characterization. All four identified viruses, one known and three new, have RNA genomes, either single-stranded (of positive or negative polarity) or double-stranded (**Table 2**). We could not identify sequences coding for polymerases among contigs belonging to the other two putative viruses, that we named *Ae. japonicus* bunyavirus 2 (AejapBV2) and *Ae. japonicus* rhabdovirus 1 (AejapRV1) based on the closest reference (**Table 2**, **Suppl. Fig. 1B, 1D**).

AejapBV2 contigs clustered with AejapTV1 based on co-occurrence in the four libraries (**Fig. 2**), but there is no further evidence that associates these sequences to the same virus. Instead, our overall analysis including small RNA size profiles (**Fig. 2**) and comparison of sequence similarity indicate that AejapBV2 and AejapTV1 are distinct viruses. Although no contigs corresponding to a putative segment L belonging to AejapBV2 were found in our libraries, we successfully assembled entire M and S genomic segments (**Suppl. Fig. 1B**). The phylogenetic analysis of amino acid sequences shows that AejapBV2 segments coding for the glycoprotein and nucleocapsid are distant from AejapBV1 equivalent sequences (**Fig. 4**). Indeed, sequence similarity and phylogenetic analysis of nucleocapsid and glycoprotein of AejapBV2 indicate that this virus belongs to the family *Phasmaviridae* in contrast to AejapBV1 that belongs to the *Phenuiviridae* family (**Fig. 4**; **Table 2**). In an attempt to find sequences encoding an RdRp belonging to AejapBV2 in our libraries, we mapped reads from each library to the segment L sequence of the virus whose segments M and S were most similar to AejapBV2 (**Table 2**), Wuhan mosquito virus 2 (GenBank: NC_031312.1) (**Suppl. Fig. 2**). Even allowing multiple mismatches, we did not observe an expressive amount of reads nor a continuous coverage of the segment L of this virus.

**Figure 4.**
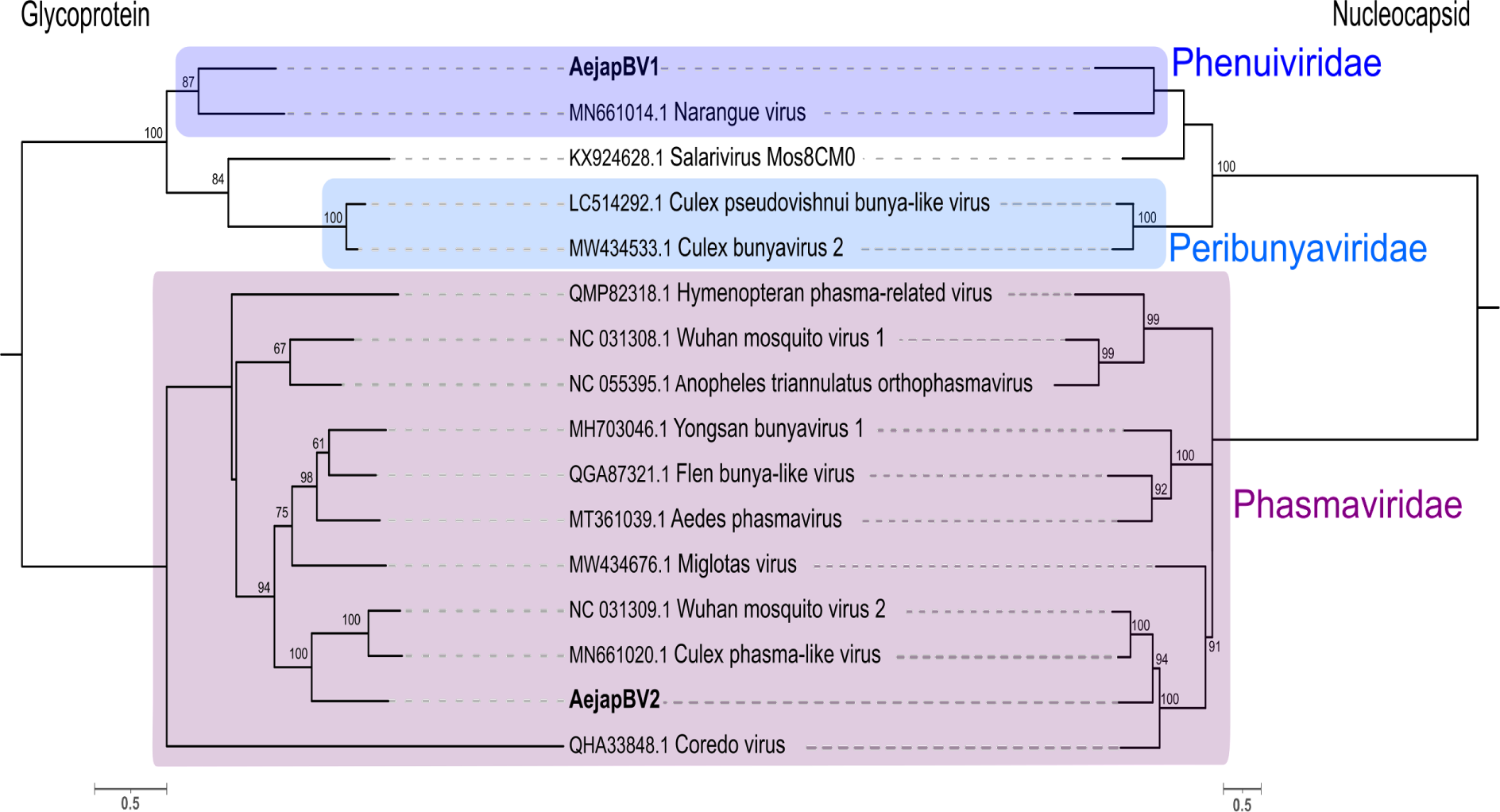
Phylogeny of glycoprotein and nucleocapsid of AejapBV1 and AejapBV2. Phylogenetic trees were generated using the glycoprotein and nucleocapsid amino acid sequences. The trees were inferred by using the Maximum Likelihood method. The trees with the highest log likelihood are shown. Number of conserved sites and the substitution models used for each tree: glycoprotein, 796 sites, WAG+G+F, and nucleocapsid, 573 sites, LG+G. Node bootstraps were calculated with 1000 replicates and are shown close to each clade and values < 60 were omitted. Trees were midpoint-rooted, and sequences from distinct viral families were included in the alignments. The trees are drawn to scale, branch lengths represent expected numbers of substitutions per amino acid site. Accession numbers for the nucleotide sequences from which the corresponding protein sequences were derived or the direct protein sequences are shown with the virus names. Viruses identified in this study are in bold.

To further characterize the identified viruses (**Table 2**), the (di)nucleotide usage of their genomes was analyzed, as in particular CpG dinucleotide usage and GC content markedly differ between the genomes of distinct virus families [61–63]. The largest representative sequence of each virus or viral segment identified was analyzed (**Table 2**). The 22 contigs of unknown origin larger than 500 nt were also included to evaluate possible associations with the viruses found in our datasets. The L, M and S segments of AejapBV1 (blue) and the M and S segments of AejapBV2 (purple) showed CpG underrepresentation and a relatively low GC content (**Fig. 5**). AejapTV1 (yellow) greatly differed from these bunyaviruses in its CpG usage and GC content (**Fig. 5**), adding yet another piece of evidence at sequence level that, despite clustering together based on co-occurrence (**Fig. 2**), AejapTV1 and AejapBV2 are indeed different viruses. The positive-stranded AejapNV1 (red) had a higher GC content compared to the negative-stranded AejapBV1, AejapBV2, AejapAV1 (green) and AejapRV1 (brown) (**Fig. 5**). This is in accordance with a previous study in which positive-sense RNA viruses were shown to have significantly higher GC contents as compared to negative-sense RNA viruses [63]. Interestingly, one of the unknown contigs (NL_02_Contig12989_12988; indicated by an arrow in **Fig. 5**) showed a high GC content and an unbiased CpG frequency, similar to AejapNV1 and narnaviruses previously discovered in other mosquito species (**Fig. 5**). As shown before, this contig clustered together with AejapNV1 based on co-occurrence and small RNA profile (**Fig. 2**; in bold). These observations reinforce the idea that this unknown contig belongs to AejapNV1.

**Figure 5.**
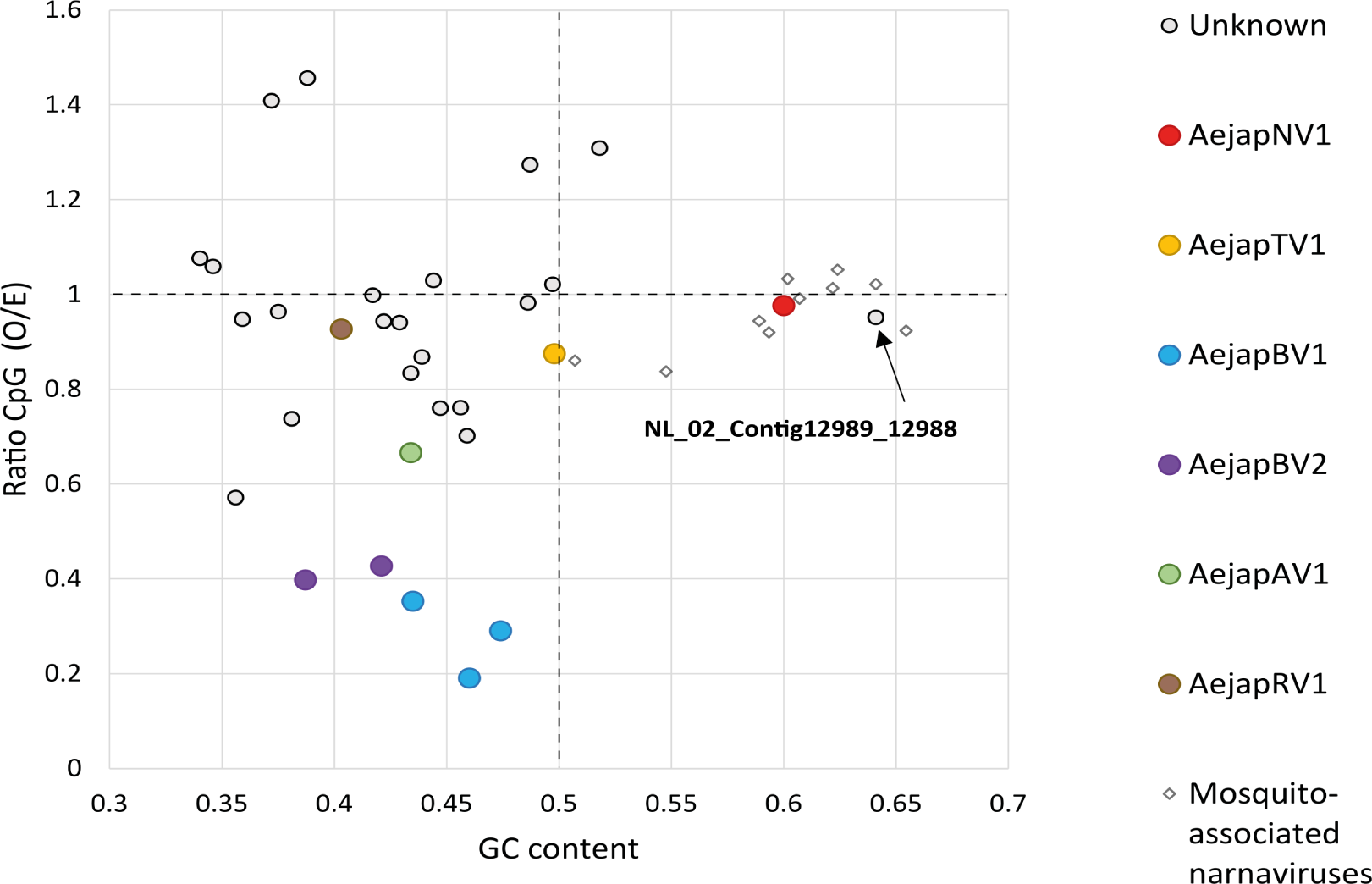
CpG dinucleotide usage and GC content of viral and unknown sequences discovered in *Ae. japonicus*. Data points indicate the ratio of GC content (X-axis) and CpG dinucleotide frequency (Y-axis) of individual viral or unknown sequences. The GC content of 0.5 (vertical dotted line) is the expected frequency if the genome would consist of 50% GC and 50% AT. The observed/expected (O/E) CpG ratio of 1.0 (horizontal dotted line) is the expected frequency of CpG occurrence when all mononucleotides in a given RNA sequence would be randomly distributed. Besides the viral and unknown sequences obtained from *Ae. japonicus*, the genome sequences of previously discovered narnaviruses in other mosquito species were also included in the analysis. GenBank accession numbers of these mosquito-associated narnaviruses are: MW226855.1, MW226856.1, NC_035120.1, KF298284.2, MF176344.1, KF298275.2, MW520409.1, MK285331.1, MK285333.1 and MK285336.1.

### Small RNA profiles and genome organisation of AejapNV1

The small RNA size profile, small RNA coverage, and ambigrammatic coding strategy of the ∼1 kb long unknown contig (NL_02_Contig12989_12988) were very similar to the ∼3 kb long AejapNV1 genome sequence coding for the RdRp (**Fig. 6A, 6B**). Together with the similarities noted before, these observations indicate that this contig is part of AejapNV1 and was thus named AejapNV1 segment 2 (S2), while the sequence encoding the RdRp is now referred to as segment 1 (S1) (**Table 2**). For S1, we observed a strong bias of mapped 24-29 nt small RNAs for the positive RdRp strand (**Fig. 6A**). Likewise, S2 also showed preference of mapped 24-29 nt small RNAs towards one specific strand (**Fig. 6A**). Analysis of the small RNA coverage profiles of AejapNV1 S1 and S2 for each small RNA length separately indicated a bias of small RNAs (18-35 nt in length) towards the same specific strand for each small RNA length, except for 21 nt small RNAs, which were found in similar quantities on both strands (**Suppl. Fig. 3**). This asymmetric pattern for small RNAs sized 18-20 nt and 22-35 nt is likely caused by non-specific degradation of viral RNAs, indicating the relative abundance and exposure of each strand [64]. For AejapNV1 S1, the small RNA bias was indeed towards the strand encoding the RdRp, which is considered the positive strand (**Suppl. Fig. 3**). Therefore we propose that the positive strand of AejapNV1 S2 is the one to which most small RNAs mapped.

**Figure 6.**
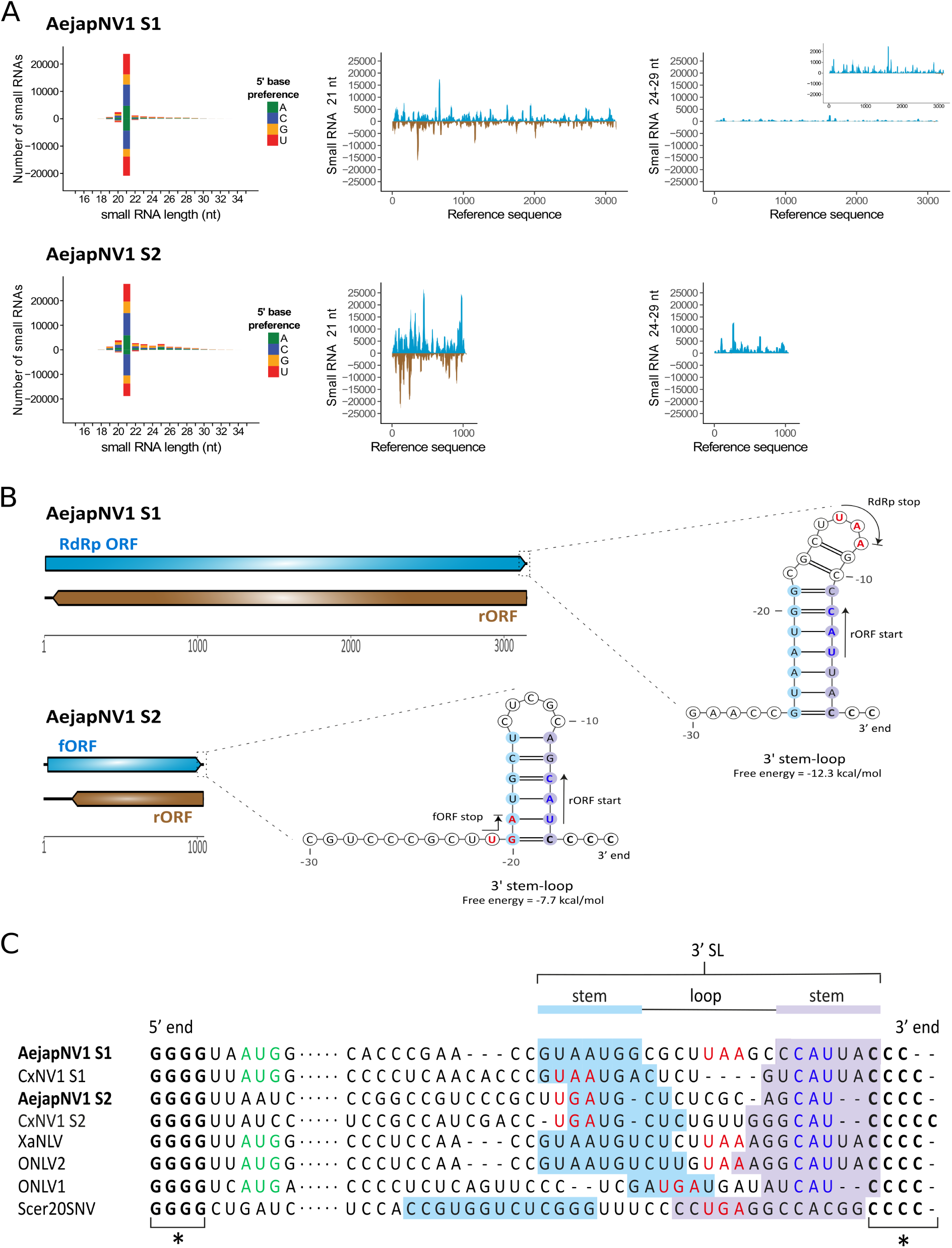
Small RNA profiles and genome organisation of AejapNV1. **(A)** Left: size distribution and 5’ base preference of small RNAs derived from AejapNV1 S1 and S2. Middle and right: coverage of 21 and 24-29 nt sized small RNAs across S1 and S2. Viral reads mapping to the positive strand are shown in blue, whereas viral reads mapping to the negative strand are shown in brown. **(B)** Genome organisation of AejapNV1. The ambigrammatic coding strategy of S1 and S2 is shown. RdRp ORF and forward ORF (fORF) on the positive strand are shown in blue, whereas reverse ORFs (rORFs) on the negative strand are shown in brown. Untranslated regions are indicated by black lines. Predicted stem-loop structures at the 3’ terminus of the positive-sense RNA strand are also shown for both segments. The locations of start and stop codons are indicated by arrows and colored blue and red, respectively. **(C)** Multiple sequence alignment of the 5’ and 3’ termini of the positive strand for indicated narnaviruses. Asterisks (*) indicate conserved, complementary runs of G or C nucleotides at the 5’ or 3’ end, respectively. Dots represent the remainder of the viral genome. Start codons of RdRp ORF are in green, stop codons of RdRp ORF / fORF are in red, start codons of rORF are in blue. The start codons of the fORFs of AejapNV1 S2 and CxNV1 S2, as well as the start codon of the RdRp ORF of Scer20SNV, are located more than 10 nt downstream of the 5’ end and therefore not shown in the alignment. The nucleotides involved in 3’ stem-loop (3’ SL) formation are indicated.

To further investigate whether the putative S2 of AejapNV1 was indeed related to S1, the 5’ and 3’ termini of both segments were analyzed for the presence of conserved narnaviral motifs. The *Alphanarnavirus* clade contains the prototypical narnaviruses such as *Saccharomyces cerevisiae* 20S RNA narnavirus (Scer20SNV) and the mosquito-associated narnaviruses containing an rORF. These viruses contain short, complementary runs of G and C nucleotides at their 5’ and 3’ termini, respectively, and have a conserved RNA stem-loop (SL) structure at their 3’ end [25, 65]. In addition, it has very recently been described that the mosquito-associated, ambigrammatic *Culex* narnavirus 1 (CxNV1) [66] also consists of two segments (an RdRp segment, hereafter named S1, and a ‘Robin’ segment, hereafter named S2), which both show these typical conserved features at the 5’ and 3’ genomic termini [67].

Using a multiple sequence alignment, the terminal sequences of AejapNV1 S1 and S2 were compared with genomes from four ambigrammatic narnaviruses found in mosquitoes (CxNV1 S1, GenBank MW226855.1; CxNV1 S2, Genbank MW226856.1; Xanthi narna-like virus (XaNLV), GenBank MW520409.1; *Ochlerotatus*-associated narna-like virus 2 (ONLV2), GenBank KF298284.2; *Ochlerotatus*-associated narna-like virus 1 (ONLV1), GenBank KF298275.1) and the yeast-infecting Scer20SNV (GenBank NC_004051.1). For all viruses and segments analyzed, short runs of G and C nucleotides were present at the 5’ and 3’ termini (**Fig. 6C**, indicated by asterisks). Based on RNA structure modelling, a conserved SL structure was predicted to occur at the 3’ terminus of AejapNV1 S1 and S2 (**Fig. 6B, 6C**). Similar conserved SL structures at the 3’ terminus, differing in size and with covarying base pairs in the stem region (**Fig. 6C**), have previously been observed for CxNV1 S1 and S2 [67], ONLV1, ONLV2 [25] and Scer20SNV [65], and were also found for XaNLV (**Suppl. Fig. 4**). The presence of these conserved narnaviral characteristics in the genomic termini of both S1 and S2 of AejapNV1 not only indicated that our small RNA sequencing method was able to recover (near) complete genomic sequences, but also corroborated the association between AejapNV1 S1 and the newly discovered S2.

### Insights into AejapNV1 S2 ORFs

To the best of our knowledge, the sequence of CxNV1 S2 is the only publicly available second segment sequence of a narnavirus infecting mosquitoes. Despite the similar genomic termini and ambigrammatic coding strategy (**Fig. 6**) [67], the S2 segments of AejapNV1 and CxNV1 are very divergent, presenting ∼46% of global identity at the nucleotide level. We therefore compared the putative polypeptides encoded by the S2 segment of these two viruses. Both forward ORFs (fORFs) and rORFs presented similar biochemical properties with high isoelectric points, suggesting a basic nature of these proteins (**Suppl. Table 6**). At the amino acid sequence level, we observed 29.34% identity for fORFs and 26.89% for rORFs (**Suppl. Fig. 5**). The predictions of secondary structures showed structured and disordered regions for the fORF encoded proteins of both viruses, with the presence of many predicted α-helices in potentially homologous regions (**Suppl. Fig. 5A**). The same pattern was not observed for the rORFs (**Suppl. Fig. 5B**). Possible transmembrane regions were detected for both fORFs within AejapNV1 S2 residues 149 to 164 and CxNV1 S2 residues 99 to 114 (**Suppl. Fig. 5A**). Using AlphaFold, we could not obtain highly confident tertiary structure predictions (pLDDT values > 90) for the entire proteins generated from the fORFs and rORFs of both viruses (**Suppl. Fig. 6**). At pLDDT values > 70, AlphaFold predicted a core structured region composed of α-helices shared by the fORFs of both viruses (**Suppl. Fig. 6A, 6C**), supporting the predicted organization of secondary structures shown in **Suppl. Fig. 5A**.

### Small RNA profiles and genome organisation of AejapTV1

Small RNAs derived from AejapTV1 (**Fig. 2**; in yellow) showed a sharp 21 nt peak matching both strands (**Fig. 7A**), which suggests activation of the siRNA pathway and active RNA replication in mosquitoes. Small RNAs mapped along the entire assembled sequence (**Fig. 7A**), which contained a capsid ORF followed by an RdRp ORF (**Fig. 7B**). Interestingly, these two ORFs were encoded in different frames (**Fig. 7B**). Ribosomal frameshifting is a strategy commonly employed by members of the family *Totiviridae* during translation of their RdRp [68, 69]. Therefore, it was investigated whether this strategy could potentially be employed by AejapTV1. Based on RNA structure modelling, a putative −1 ribosomal frameshift area was discovered at the end of the capsid ORF (**Fig. 7B**). A slippery site (GGAAAAC), present just before the stop codon of the capsid ORF, corresponded to the heptameric consensus motif typical for −1 ribosomal frameshifts and represents the area where the ribosome shifts back into another reading frame [70]. Right after the slippery site, a spacer region of 5 nucleotides in length was found (**Fig. 7B**). This region was followed by a highly structured area consisting of a three-stemmed pseudoknot (**Fig. 7B**), which is expected to be responsible for pausing and relocating the ribosome. Similarly, a −1 ribosomal frameshift has also been predicted at the end of the capsid ORF for the mosquito-associated totiviruses *Armigeres subalbatus* totivirus and Omono River virus [71–73], suggesting that −1 ribosomal frameshifting is a common feature of these mosquito-associated totiviruses.

**Figure 7.**
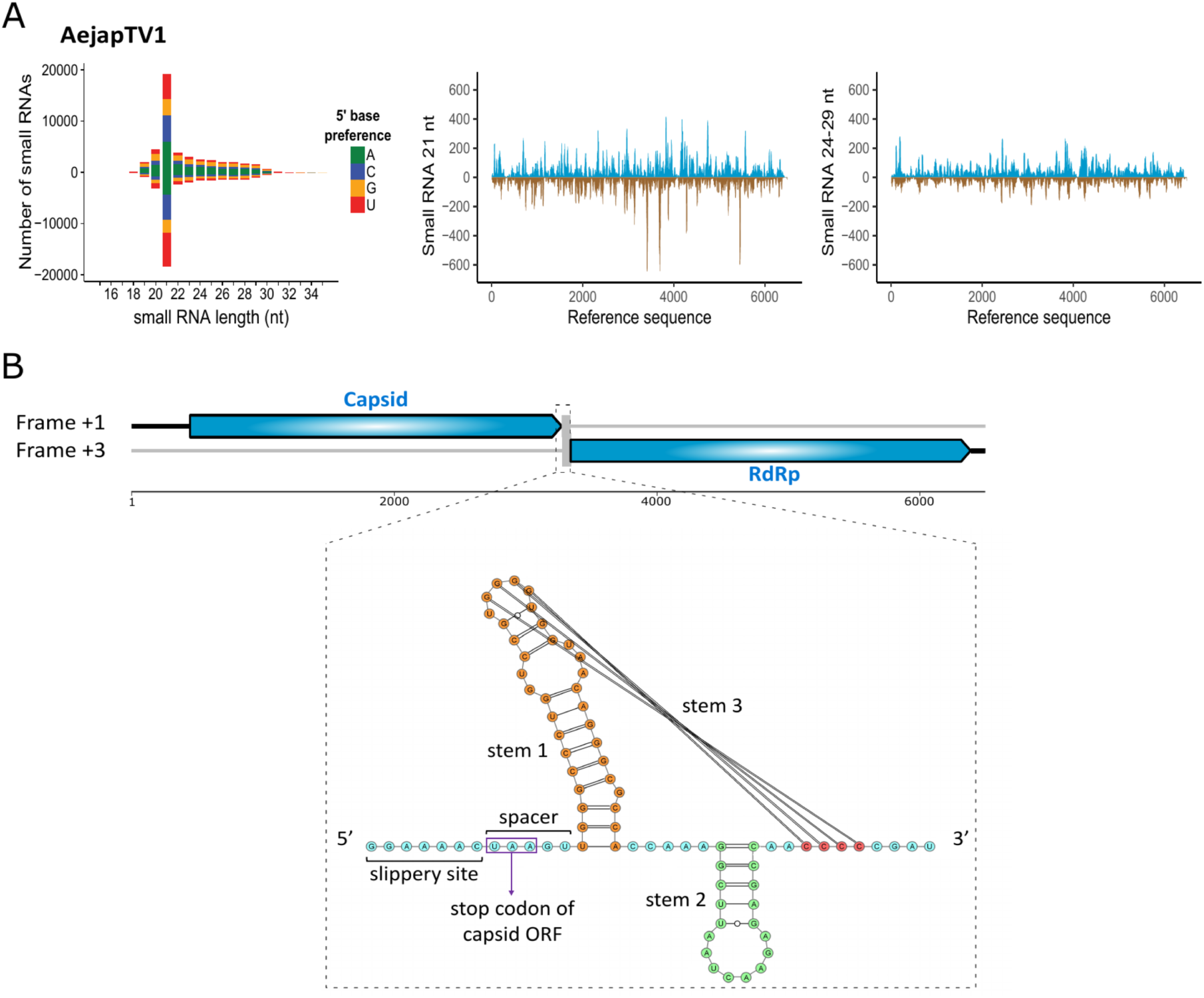
Small RNA profiles and genome organisation of AejapTV1. **(A)** Left: size distribution and 5’ base preference of small RNAs derived from AejapTV1. Middle and right: coverage of 21 and 24-29 nt sized small RNAs across the genome of AejapTV1. Viral reads mapping to the positive strand are shown in blue, whereas viral reads mapping to the negative strand are shown in brown. **(B)** Genome organisation of AejapTV1. Untranslated regions are indicated by black lines. Capsid and RdRp ORFs on the positive strand are shown in blue. These ORFs are encoded in different frames, and a putative −1 ribosomal frameshift area was observed in between the two ORFs. This area consisted of a slippery heptamer, a spacer region, and a predicted three-stemmed pseudoknot with a free energy of −33.97 kcal/mol.

### Small RNA profiles of AejapBV1, AejapBV2, AejapAV1 and AejapRV1

For the two bunyaviruses in our datasets, AejapBV1 and AejapBV2, a peak consisting of 21 nt small RNAs (characteristic of siRNA pathway activation) was observed for all three (**Fig. 8A**) or two (**Fig. 8B**) segments, respectively, and siRNAs aligned across entire segments. Also, RNAs with characteristics of piRNAs (∼24-29 nt in length) were detected for both viruses (**Fig. 8A, 8B**). Those RNAs showed U enrichment at the 1st nt of antisense reads and A enrichment at the 10th nt of sense reads, further suggesting their piRNA biogenesis (**Fig. 8A, 8B**). Moreover, reads that mapped to the segments M and S from both viruses presented a significant 10 nt overlap between small RNAs from sense and antisense strands, which is in accordance with the piRNA signature of the ping-pong amplification mechanism [74]. In a similar fashion to AejapBV1 and AejapBV2, small RNAs derived from AejapAV1 and AejapRV1 also showed a 21 nt siRNA peak, as well as ∼24-29 nt sized piRNAs with ping-pong amplification features (**Fig. 9A, 9B**). In the case of AejapRV1, this pattern reinforces the idea that we identified a real exogenous rhabdovirus that is replicating in *Ae. japonicus* rather than an EVE in the mosquito genome, despite the absence of an RdRp sequence.

**Figure 8.**
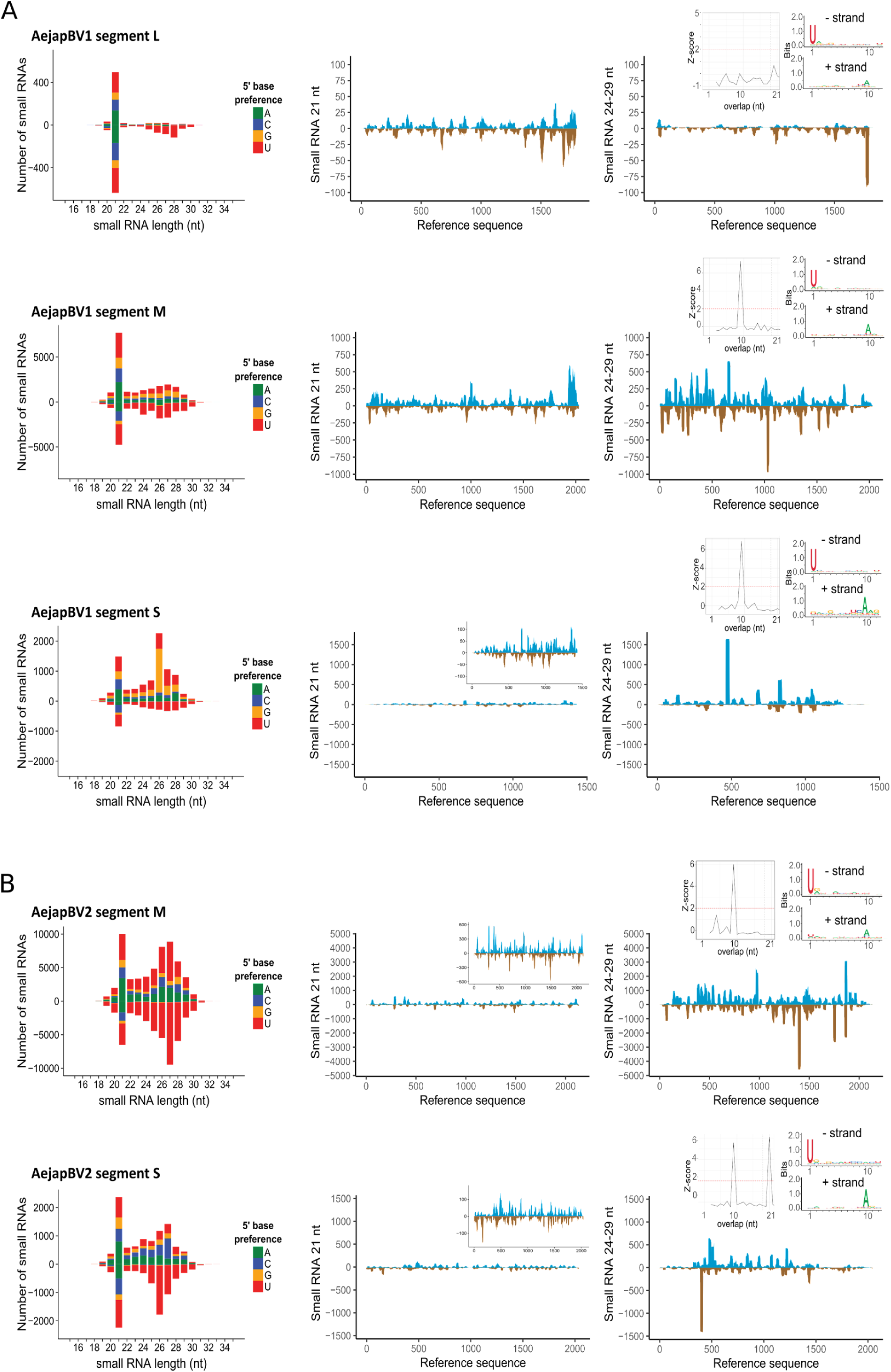
Small RNA profiles of AejapBV1 and AejapBV2. Size distribution and 5’ base preference of small RNAs derived from **(A)** AejapBV1 segments L, M and S and **(B)** AejapBV2 segments M and S are shown on the left, whereas coverage of 21 and 24-29 nt sized small RNAs across the same segments of the respective viruses is shown in the middle and on the right. For the coverage profiles, viral reads mapping to the positive strand are indicated in blue, whereas viral reads mapping to the negative strand are indicated in brown. Inset figures in the middle figures show a zoomed in 21 nt small RNA coverage. Inset figures in the right figures show features of 24-29 nt reads aligned to its respective viral references. Left inset figures show a histogram with normalized frequency of overlap sizes of reads aligned to positive and negative strands. Right inset figures show sequence logo representations of reads aligned to positive and negative strand separately.

**Figure 9.**
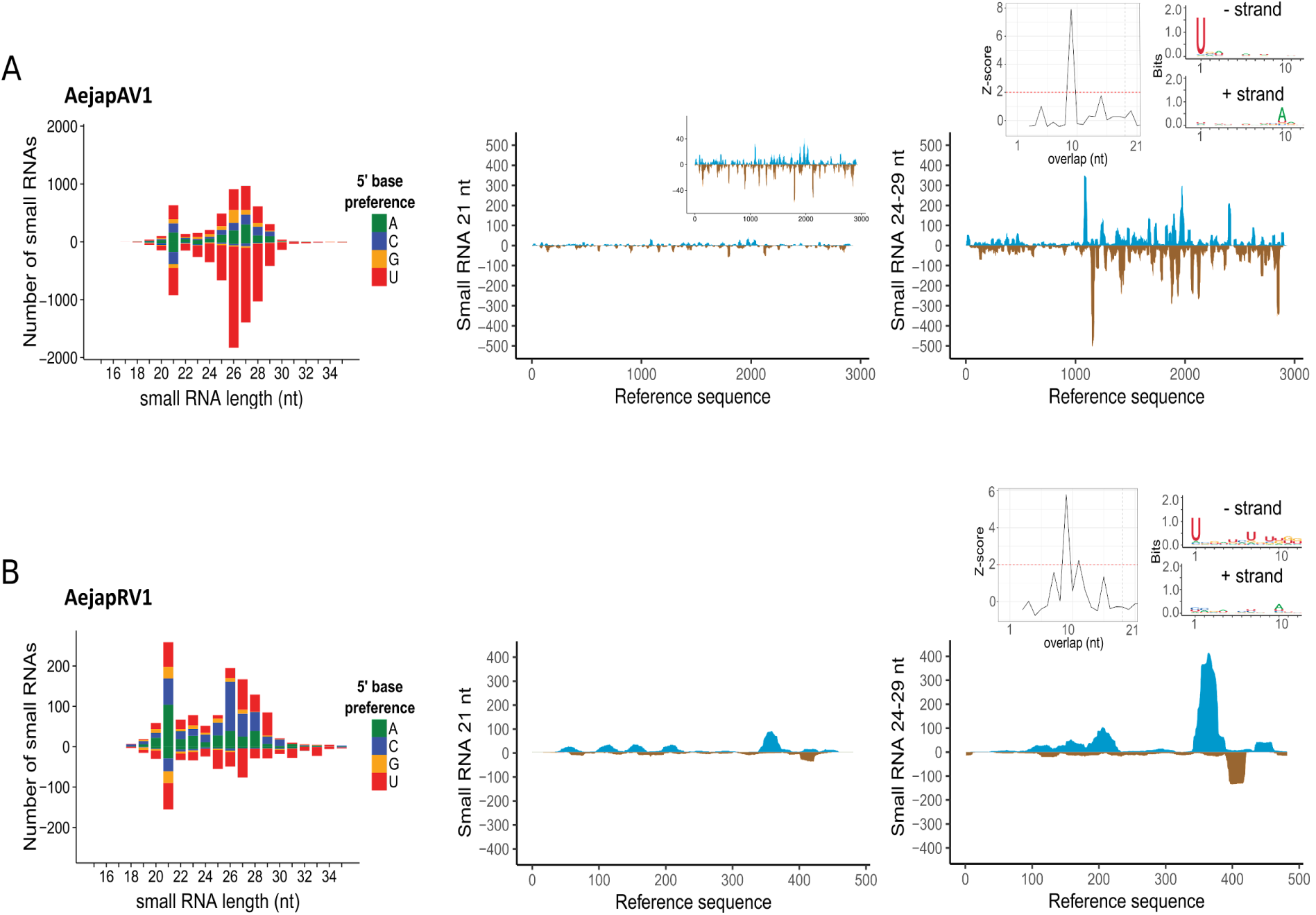
Small RNA profiles of AejapAV1 and AejapRV1. Size distribution and 5’ base preference of small RNAs derived from **(A)** AejapAV1 and **(B)** AejapRV1 are shown on the left. In the middle and on the right, the coverage of 21 and 24-29 nt sized small RNAs across the assembled contigs of the same respective viruses is shown. For the coverage profiles, viral reads mapping to the positive strand are indicated in blue, whereas viral reads mapping to the negative strand are indicated in brown. Inset figure in middle figure A shows a zoomed in 21 nt small RNA coverage. Inset figures in the right figures show features of 24-29 nt reads aligned to its respective viral references. Left inset figures show a histogram with normalized frequency of overlap sizes of reads aligned to positive and negative strands. Right inset figures show sequence logo representations of reads aligned to positive and negative strand separately.

### Prevalence of viruses in field-collected *Ae. japonicus* mosquitoes

To confirm whether the *de novo* assembled viruses were present in field-collected *Ae. japonicus*, RT-PCR assays were designed using primers targeting the RdRp coding region of AejapNV1, AejapTV1, AejapAV1 or AejapBV1. RT-PCRs with primers targeting S2 of AejapNV1 were also performed to confirm the presence of this putative segment. All four viruses were successfully detected in (pools of) field-collected adult *Ae. japonicus* females, whilst they could not be detected in adult *Cx. pipiens* females that were tested in parallel (**Fig. 10A**). For AejapNV1, the presence of both S1 and S2 in *Ae. japonicus* was confirmed (**Fig. 10A**). DNA sequencing of the AejapNV1 S2 RT-PCR product showed 99.6% identity to the contig of AejapNV1 S2 previously obtained from *de novo* small RNA assembly. These analyses confirm that our metagenomic approach successfully identified viruses and did not generate artifacts of assembly.

**Figure 10.**
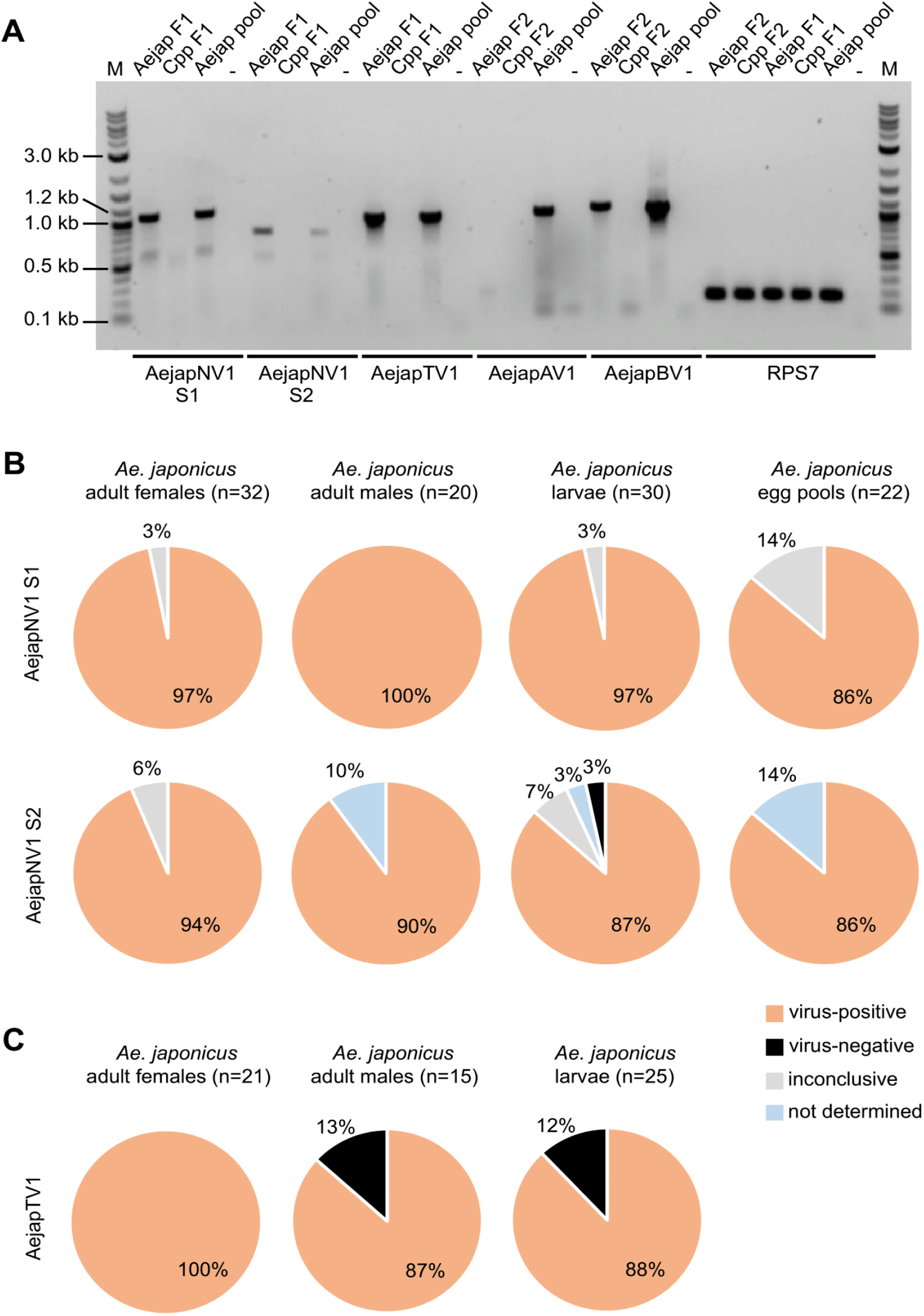
Detection of AejapNV1 S1 and S2, AejapTV1, AejapAV1 and AejapBV1 in field-collected *Ae. japonicus* from the Netherlands by RT-PCR. **(A)** Individual adult female *Ae. japonicus* (Aejap F1, F2), individual adult female *Cx. pipiens pipiens* (Cpp F1, F2), a pool of four *Ae. japonicus* adult females (indicated by ‘Aejap pool’) and a water sample (no RNA; negative control, indicated by ‘-’) were tested for presence of the viruses. All samples were also tested for mosquito ribosomal protein S7 (RPS7) to check the quality of the RNA. Expected amplicon sizes were: 1060 bp (AejapNV1 S1), 774 bp (AejapNV1 S2), 998 bp (AejapTV1), 1043 bp (AejapAV1), 1095 bp (AejapBV1) and 175 bp (RPS7). Aejap F1 tested positive for AejapNV1 S1 and S2, and also for AejapTV1. Aejap F2 tested negative for AejapAV1, but positive for AejapBV1. The pool of *Ae. japonicus* females was positive for all tested viruses. *Cx. pipiens* females tested negative for all viruses. The lanes indicated with ‘M’ contain the DNA marker. **(B)** Prevalence of AejapNV1 S1 and S2 in *Ae. japonicus* adult females, adult males, larvae and pools of 25 eggs. Individual samples were screened by RT-PCR. The number of samples tested is indicated by ‘n’. Results were marked as inconclusive when samples tested virus-negative the first time, and this could not be confirmed a second time due to lack of RNA, whereas parallel samples in the same assay which tested virus-negative the first time, tested virus-positive the second time. **(C)** Prevalence of AejapTV1 in *Ae. japonicus* adult females, adult males and larvae. Individual samples were screened by RT-PCR. The number of samples tested is indicated by ‘n’.

Next, *Ae. japonicus* adult females, adult males, larvae and egg pools were screened for the presence of AejapNV1 S1 and S2. Both segments were present at very high prevalence (close to 100%) in all tested mosquito life stages (**Fig. 10B**). Screening of *Ae. japonicus* mosquitoes for AejapTV1 resulted in 100% virus-positive adult females, 87% virus-positive adult males and 88% virus-positive larvae (**Fig. 10C**).

## DISCUSSION

In this study, we analysed the virome of field-collected *Ae. japonicus* and found four viruses, three of them novel species, and another two putative novel viruses associated with this invasive vector mosquito. We applied a small RNA-based metagenomic approach previously developed by our group [28] that allowed us to overcome three main challenges of eukaryotic viral metagenomics: i) the differentiation of replicating exogenous viruses from EVEs based on the presence of siRNAs, ii) the association of different segments of the same virus based on co-occurrence and small RNA profiles, and iii) the identification and classification of highly divergent viral sequences that do not have similarity to any known reference sequence.

All viruses detected in *Ae. japonicus* have RNA genomes [12, 23]. Detection of DNA viruses using RNA-seq approaches is not supposed to be a limitation since DNA viruses produce different RNA molecules during their replication cycle [75]. Indeed, DNA viruses have been successfully identified in mosquitoes via metatranscriptomics approaches based on long and small RNA sequencing [76, 77]. From a broader perspective, DNA viruses seem to be underrepresented in mosquito viromes [15]. Therefore, DNA viruses may indeed be absent in the *Ae. japonicus* virome rather than undetected due to methodological limitations.

Regarding our results on the identification of *Ae. japonicus*-associated viruses, AejapNV1 is a good example of the power of our strategy. *De novo* assembly of small RNAs did not only yield the ∼3 kb long primary genome segment for AejapNV1, but interestingly also revealed the presence of a ∼1 kb long secondary genome segment (also ambigrammatic, but containing ORFs with no hits in the databases), which we could confidently associate to the same virus. Small RNA abundancy and size distribution profiles were very similar for both segments, suggesting that they belong to the same virus. Further support comes from the similar CpG and GC content, conserved genomic termini and unusual ambigrammatic nature of segments S1 and S2. Bisegmented narnaviruses have also been discovered in *Plasmodium* [78], in a trypanosomatid [79, 80], and, very recently, in mosquitoes [81, 82]. For the mosquito-associated, bisegmented, ambigrammatic CxNV1, the presence of the primary RdRp segment was found to be required for replication of the secondary (‘Robin’) segment in *Culex* mosquito cells, whereas the primary segment could persist in cell culture without the presence of the secondary segment [67]. In our study, the primary and secondary genome segment of AejapNV1 co-occurred at very high frequency in field-collected *Ae. japonicus* mosquitoes, which is in accordance with previous findings for CxNV1 in wild-caught mosquitoes [81], thus suggesting that the secondary segment is of key importance for virus survival in the field.

Despite some conserved features between AejapNV1 S2 and CxNV1 S2, such as the genomic termini and ambigrammatic ORF structure, these sequences are highly divergent at nucleotide and amino acid sequence level. Such divergence raises questions about the origin and function of this secondary genome segment. Here we analyzed the conservation between polypeptides encoded by AejapNV1 S2 and CxNV1 S2 at the structure levels. Conserved motifs at secondary and tertiary levels between proteins encoded by the fORFs were observed when comparing the presence and synteny of predicted α-helix regions and the core structure predicted by AlphaFold. Interestingly, the rORF of CxNV1 S2 may not encode a functional protein [82]. Nevertheless, the function of the secondary genome segment of AejapNV1 and CxNV1 remains unknown, and future studies are needed to elucidate its role and evolutionary origin. These narnavirus secondary segments belong to the dark matter of metagenomics, since sequence similarity methods, such as local alignments, were not able to associate the AejapNV1 S2 sequence to the CxNV1 S2 reference in GenBank. Even the application of the powerful tool AlphaFold [59] to predict tertiary structures and infer protein function, was limited by the lack of similar sequences to ORFs in AejapNV1 S2. AlphaFold relies on initial alignments to protein sequence and structure databases, such as Uniref90 and the Protein Data Bank, to acquire atomic tridimensional data and perform structural assessments [59]. This could also explain the lack of confident predictions for our fORFs and also for any attempt of predicting structures for highly divergent sequences. This is a good example of how bioinformatics pipelines for viral sequence identification in metagenomic data still have room for improvement. Increased knowledge about the viral metagenomic dark matter is paramount to allow large-scale automated generalizations of the evolution and function of divergent viral sequences. Of note, our clustering approach based on small RNA profiles and co-occurrence was critical to identify the unknown contig corresponding to segment 2 of AejapNV1. This reinforces the power of small RNA-based approaches to recover viral sequences from the metagenomic dark matter [28].

Still, the origin of a substantial portion of the dark matter present in *Ae. Japonicus* remains enigmatic. Despite the presence of an siRNA profile, we could not clearly associate 21 unknown contigs to any defined viral species in this study. Further studies will be necessary to determine if these unknown sequences with siRNA profile are of viral origin. One possible explanation derives from the lack of a reference genome available for *Ae. japonicus*, implying that some of those unknown contigs could have originated from unknown repetitive elements in the genome and/or overlapping mRNA fragments that may produce endogenous siRNAs [83].

Our small RNA-based approach also pointed to the presence of two distinct bunyaviruses in *Ae. japonicus*. However, we could not identify a potential L segment encoding an RdRp for AejapBV2. Based on previous successful detection of segmented viruses from different families including bunyaviruses using our small RNA-based approach [18, 28], it is unlikely that the segment L of AejapBV2 was present in the samples and remained undetected. In addition, we successfully assembled complete M and S segments for AejapBV2. Segment reassortments between bunyaviruses are frequently described and it has been suggested that most if not all bunyaviruses currently present have arisen from reassortments [84, 85]. Given the ability of bunyaviruses to reassort genome segments, AejapBV2 could possibly use the RdRp of AejapBV1 for replication. However, AejapBV1 and AejapBV2 belong to two different bunyavirus families, *Phenuiviridae* and *Phasmaviridae*, respectively, which might render complementation or reassortment less likely. In addition, this hypothesis implies that AejapBV2 replication is dependent on the presence of the L segment of AejapBV1, but we could not detect AejapBV1 in our *Ae. japonicus* NL_02 small RNA library whilst AejapBV2 was present (**Fig. 2**). Future studies will be needed to elucidate the replication strategy of AejapBV2 in *Ae. japonicus*.

Remarkably, AejapBV1, AejapBV2, AejapAV1 and AejapRV1 produced abundant piRNAs with strong signals of a ping-pong mechanism. Notably, not all viruses induce the production of piRNAs during infection in mosquitoes [74, 86]. A similar small RNA profile was described for another bunyavirus, Phasi Charoen-like virus, infecting *Ae. aegypti*, and the authors reasoned that the piRNA biogenesis could be linked to ovary tropism [28]. It would be of interest to investigate if the piRNA profiles observed in our study correlate with tissue tropism in *Ae. japonicus*. Interestingly, there is a striking difference in the profile of piRNAs produced from the segments M and S of AejapBV1 and AejapBV2, which suggests that these viruses infect different tissues in the mosquito [28].

Screening of wild-caught *Ae. japonicus* from the Netherlands indicated a high prevalence of AejapNV1 and AejapTV1 across all tested mosquito life stages, suggesting efficient transmission of these viruses, possibly vertically. Such a phenomenon was described for narnaviruses discovered in fungi [87] and nematodes [88], for which vertical transmission has been shown. The high percentage of mosquitoes that tested positive for AejapNV1 in our study is also in accordance with a recent study in Japan, where all tested *Ae. japonicus* mosquitoes harboured AejapNV1 [23].

In our study, AejapNV1, AejapTV1, AejapAV1, AejapBV1 and AejapBV2 were all detected in *Ae. japonicus* mosquitoes from the Netherlands and France. This indicates a high stability of the virome across mosquito populations, which has also been observed for other mosquito species [20, 22], and may suggest that these viruses have co-evolved with their mosquito host. Although the newly discovered virus species belong to different virus families and are thus highly diverse, they form a stable viral community in *Ae. japonicus* despite active antiviral RNAi responses, hence suggesting that these viruses should be considered important constituents of the biology of *Ae. japonicus* mosquito populations.

## CONCLUSIONS

This study uncovered the virome of the invasive Asian bush mosquito *Ae. japonicus*, a species that can transmit multiple arboviruses and has become increasingly prevalent in North America and Europe. We discovered a highly diverse virome in *Ae. japonicus* with viruses from at least five families: *Narnaviridae*, *Totiviridae*, *Xinmoviridae*, *Phenuiviridae* and *Phasmaviridae*. We showed a collection of viruses not only diverse at the sequence level but also in RNAi responses generated by *Ae. japonicus*, providing the basis for future studies to elucidate the immune mechanisms involved in dealing with multiple viral infections. By discovering and characterizing AejapNV1 S2, we showed the strength of our small RNA-based approach to access the “viral dark matter” of insect metagenomes. Beyond associating a highly divergent viral genomic segment to a single virus, we provided evidence that it replicates in the host by showing its siRNA signature. Our study establishes solid ground to explore whether these resident viruses could impact arbovirus transmission by *Ae. japonicus*. Infection with these likely ISVs might add yet another complex variable to the risk assessment of arbovirus outbreaks caused by *Ae. japonicus*, and future studies are needed to dissect the role of the virome in arbovirus transmission.

## DECLARATIONS

### Ethics approval and consent to participate

Not applicable.

### Consent for publication

Not applicable.

### Availability of data and material

The small RNA datasets generated and/or analysed during the current study are available in the NCBI sequence read archive (SRA) under BioProject PRJNA545039 and PRJNA785589. The genome sequences of newly discovered viruses can be found in GenBank under NCBI accession numbers OP737829-OP737837.

### Authors’ contributions

SRA, JPPA, RPO, ERGRA, GPP and JTM conceptualized the study. SRA, RPO, CB, JSG, CL, CJMK, and EM designed and conducted the field work and the wet lab experiments. SRA, JPPA, RPO, CB, JSG, BMS, JJF and ERGRA designed and performed the bioinformatic analyses. SRA, JPPA, GPP and JTM coordinated the study. SRA and JPPA wrote the draft manuscript. SRA, JPPA, RPO, CB, JSG, CL, CJMK, BMS, JJF, ERGRA, EM, GPP and JTM revised the manuscript. All authors read and approved the final manuscript.

## Acknowledgements

We thank Marleen Abma-Henkens for her assistance with *Ae. japonicus* collection in the field, Pieter Rouweler and other members of the insect rearing group from the Laboratory of Entomology from Wageningen University & Research for maintaining the *Cx. pipiens* colony, Giel Göertz for his contributions during the start of the project, Monique van Oers for her continued interest in the project, and Eric Snijder, Andrew Firth, Nina Lukhovitskaya and Katherine Brown for fruitful scientific discussions. We also thank Isaque J. S. de Faria and all members of Marques Laboratory, who contributed with suggestions and discussions and Rune Hartmann for his insights about tertiary structure predictions.

## ADDITIONAL FILES

**Additional file 1 –** Supplementary Figures

**Additional file 2 –** Supplementary Tables

Supplementary Table 1. Library information.

Supplementary Table 2. Assembly metrics.

Supplementary Table 3. Contig classification.

Supplementary Table 4. Viral BLAST hits per library.

Supplementary Table 5. CD-HIT summary.

Supplementary Table 6. Biochemical properties of proteins encoded by ORFs in AejapNV1 S2 and CxNV1 S2.

Supplementary Table 7. AlphaFold model ranking.

## Supplementary Figures

**Supplementary Figure 1.**
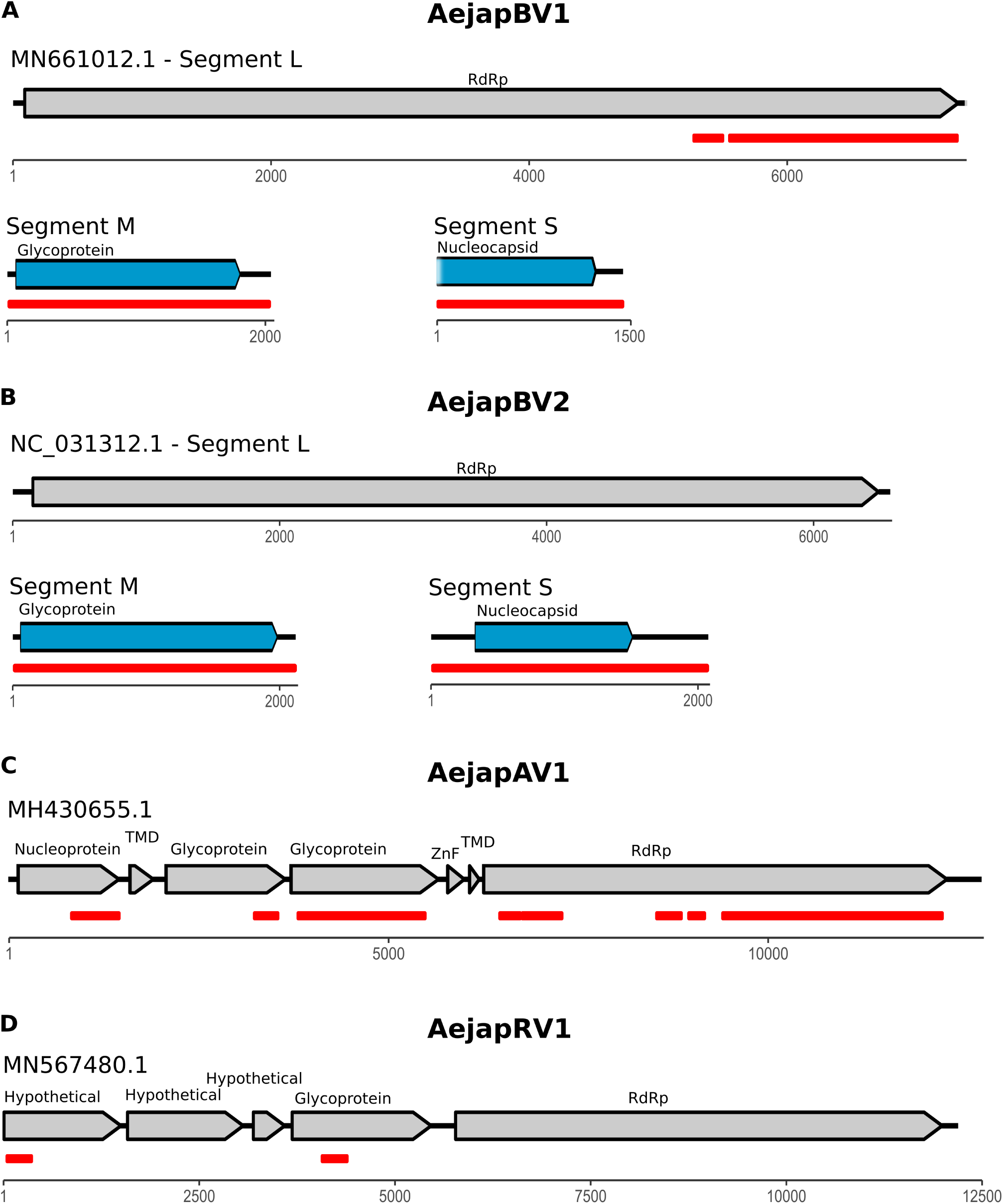
Genomic organization of partially assembled viruses. Grey arrows represent ORFs from the closest GenBank reference viral sequence. Red lines indicate viral reference genome regions covered by our assembled contigs. Blue arrows represent ORFs from completely assembled viral segments in this work. Black lines indicate untranslated regions. **(A)** AejapBV1. The lack of an assembled 5’ UTR for segment S is represented as a fading color region. Despite the lack of a 5’ UTR and a start codon, the total ORF size of segment S is similar to its closest sequence in GenBank (QHA33859.1). **(B)** AejapBV2, **(C)** AejapAV1, **(D)** AejapRV1.

**Supplementary Figure 2.**
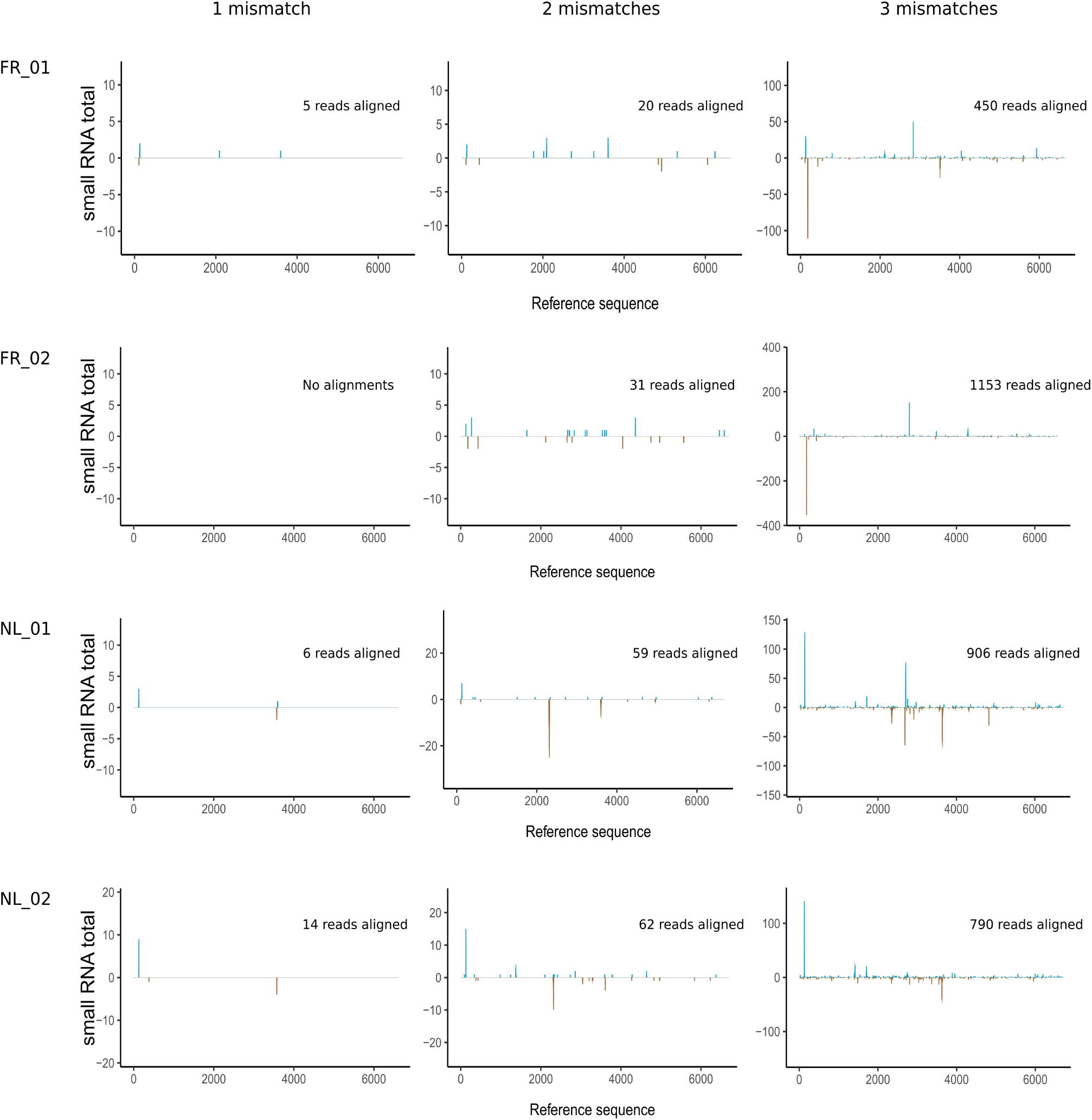
Small RNA coverage of Wuhan mosquito virus 2 segment L. Total small RNA reads from each library (rows) were aligned to Wuhan mosquito virus 2 segment L (NC_031312.1), a potential homologous sequence of the missing segment L of our identified AejapBV2. In order to confirm the absence of a highly divergent segment L inexplicably not assembled, we aligned each small RNA library allowing one to three mismatches per read (columns) by varying the parameter *-v* of bowtie. Each panel represents the coverage and total reads assembled for the combination of one library and the maximum number of mismatches allowed per read. The blue lines indicate small RNA read coverage of forward strand and the brown lines of negative strand. Only when three mismatches were allowed, considerable numbers of reads aligned to the reference. Although with no signal of continuous coverages, these results likely indicate spurious alignments.

**Supplementary Figure 3.**
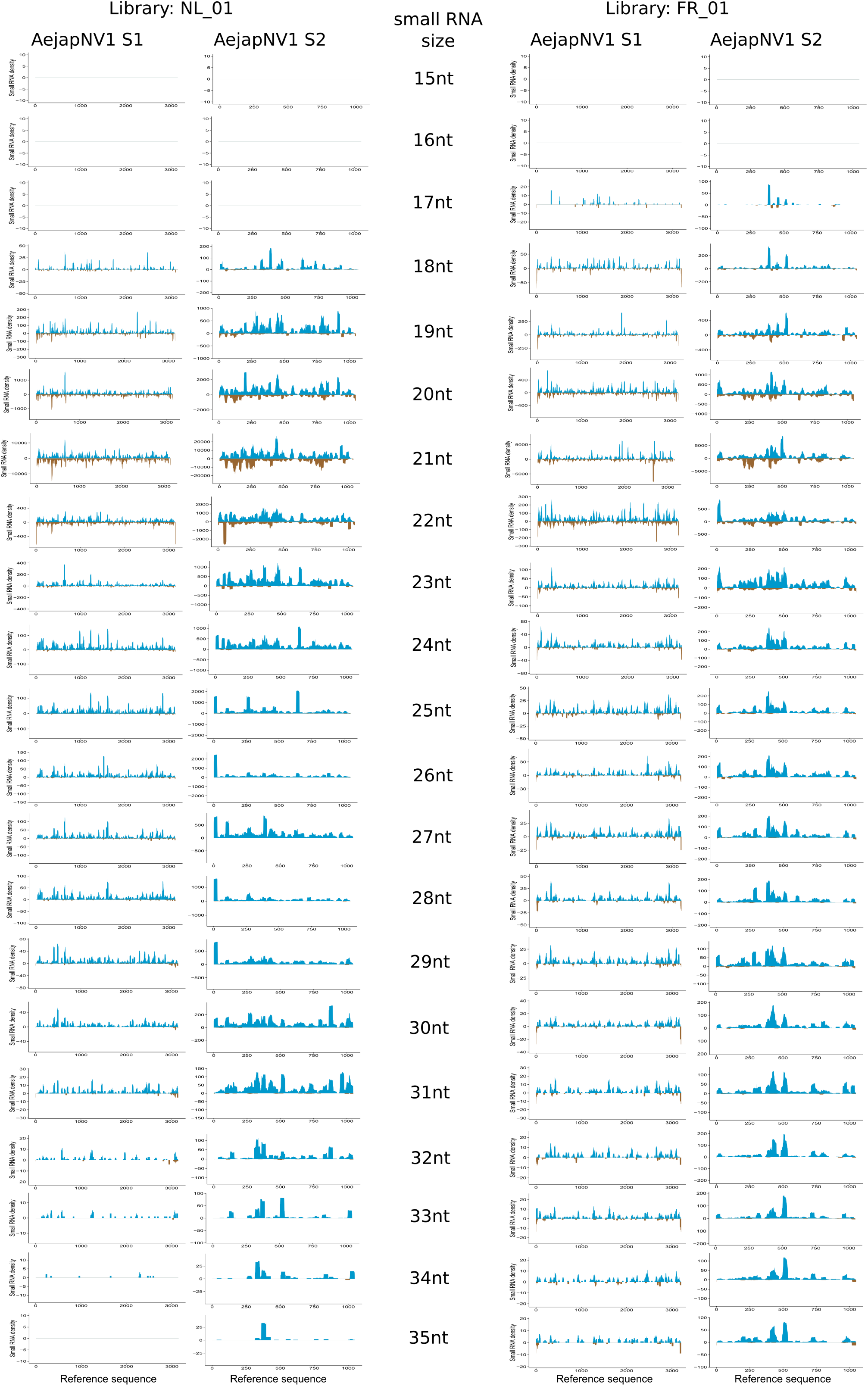
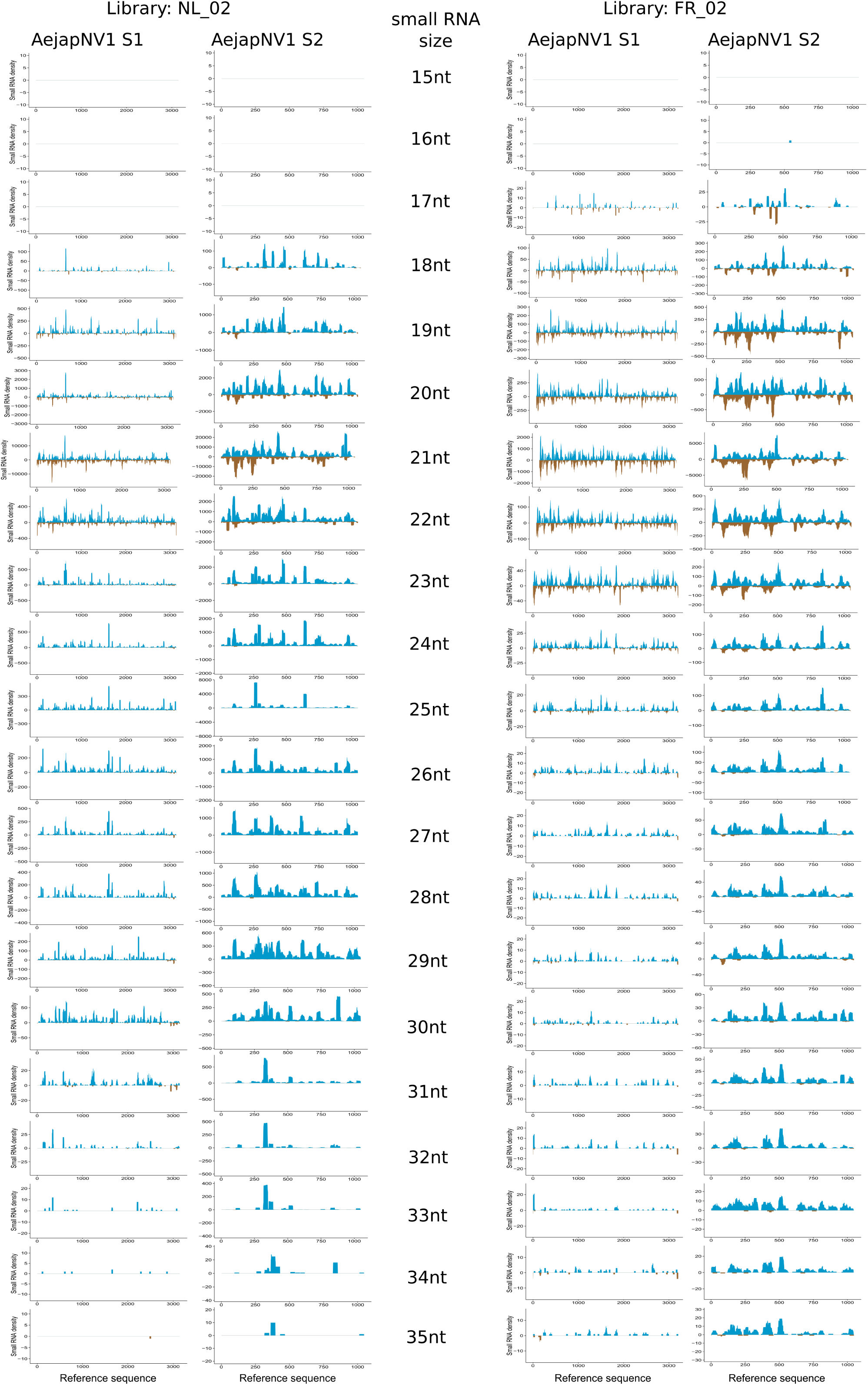
Strand-specific small RNA coverage bias of AejapNV1 51 and 52. Reads from sizes 15 to 35 nt from each library were aligned separately to segments 81 and 82. The blue area indicates small RNA read coverage of forward strand and the brown area of negative strand. The 81 sequence orientation was determined based on the RdRp coding ORF direction. A small RNA coverage bias towards the positive RdRp strand of 81 was seen, and a similar coverage bias was seen for 82, thus indicating the putative positive strand.

**Supplementary Figure 4.**
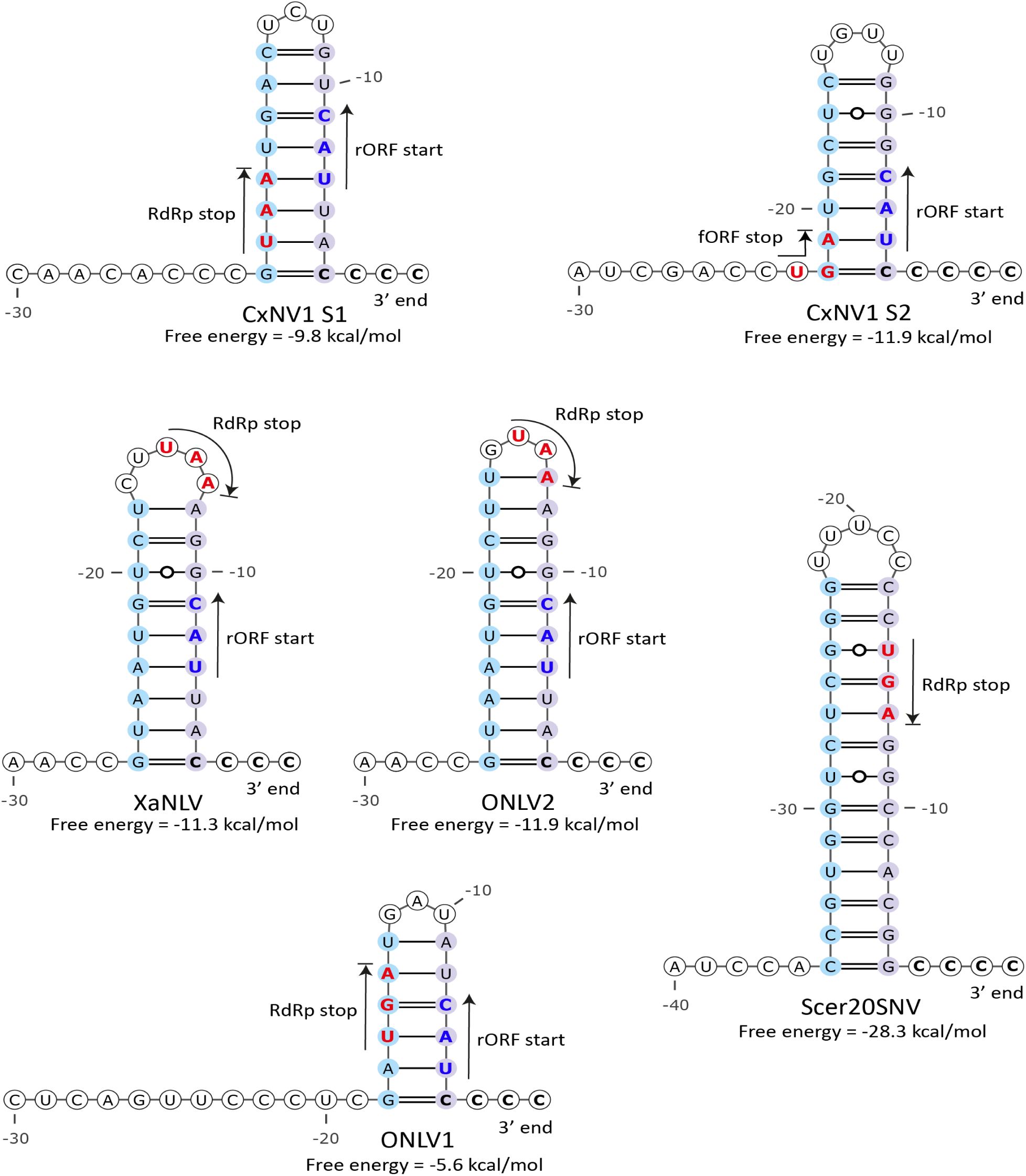
Predicted 3’ stem-loop structures of CxNV1 S1, CxNV1 S2, XaNLV, ONLV2, ONLV1 and Scer20SNV. The predicted RNA structures at the 3’ terminus of the positive-sense RNA strand are shown. Locations of start and stop codons are indicated by arrows and colored blue and red, respectively.

**Supplementary Figure 5.**
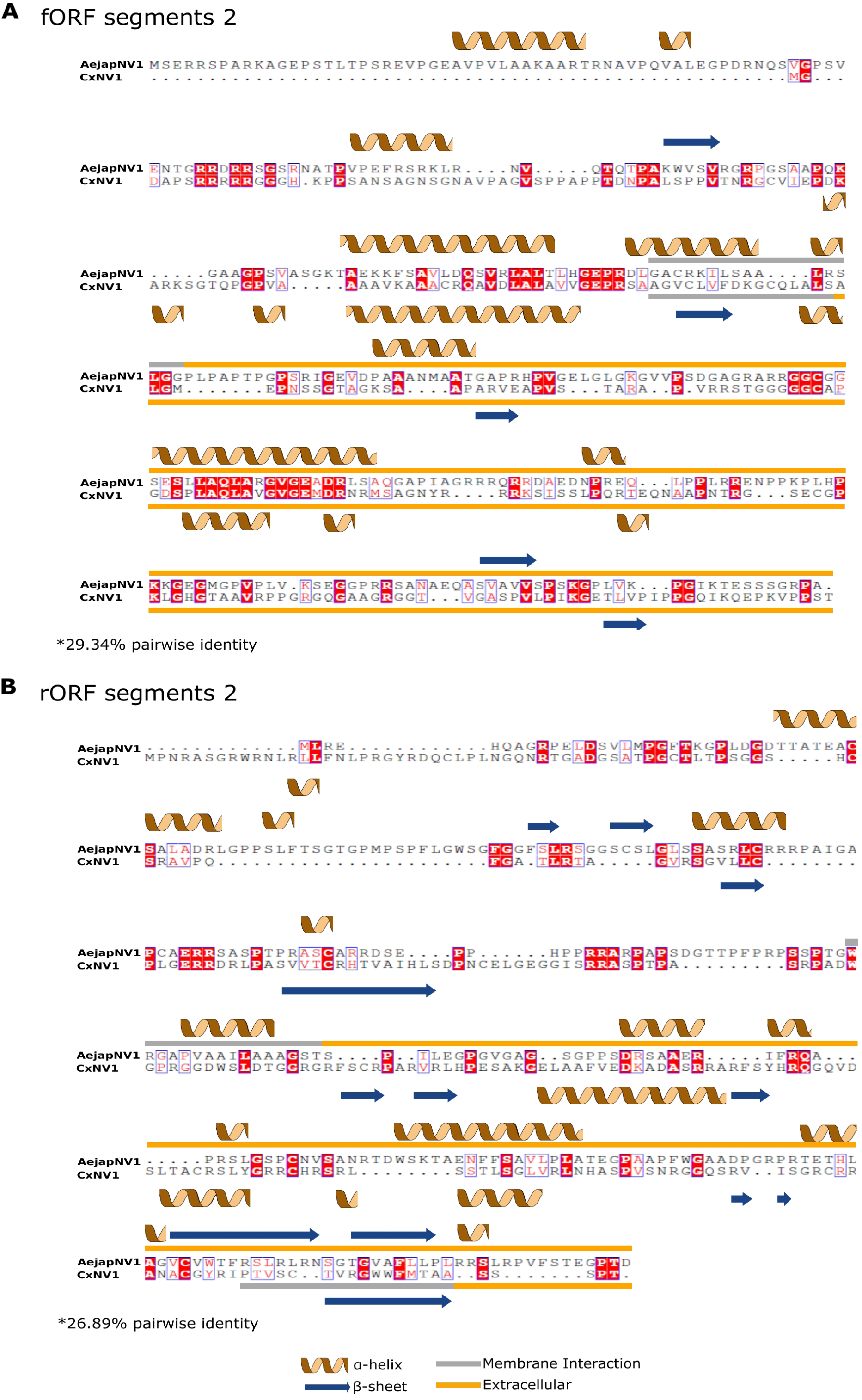
Primary and secondary protein structure comparisons of segment 2 ORFs from AejapNV1 and CxNV1. Sequence alignment of AejapNV1 and CxNV1 (GenBank MW226856.1) fORF and rORF segment 2 and its respective secondary structure prediction and membrane interaction. In **(A)** are shown the comparisons of fORFs and in **(B)** the rORFs. Residues with identical matches are colored in red boxes, and residues with similar side chain physical-chemical properties are highlighted with a blue box and written in red. Above the sequences are represented the PSIPRED and MEMSAT-SVM secondary structure prediction results of AejapNV1, and under the sequences, the CxNV1 prediction. The α-helix regions are represented with brown helices and the β-strands with dark blue arrows. The regions with predicted membrane interaction are in gray boxes and predicted extracellular regions in yellow.

**Supplementary Figure 6.**
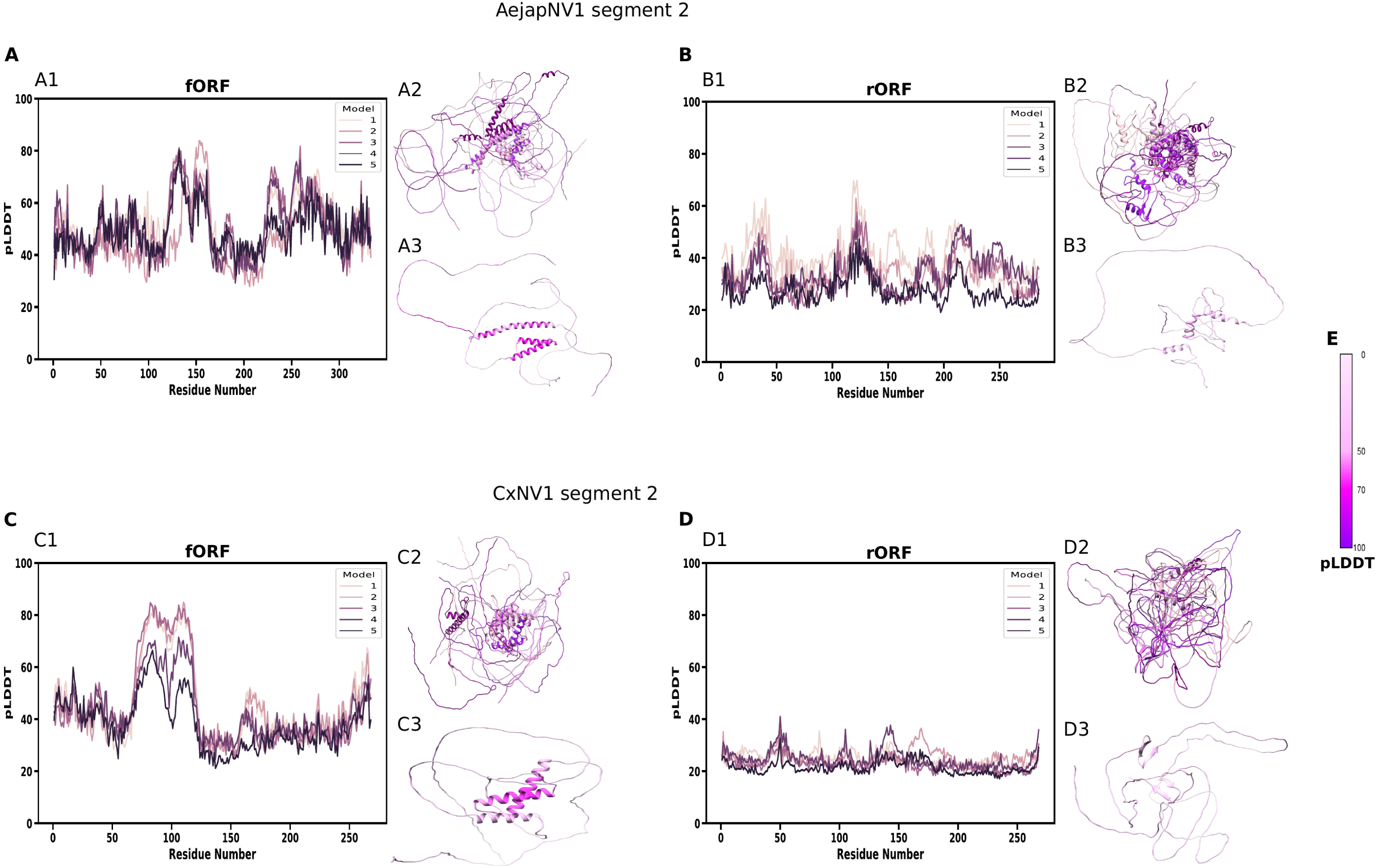
Tertiary protein structure predictions for segment 2 ORFs of AejapNV1 and CxNV1. AlphaFold per-residue confidence estimation (pLDDT) and calculated models are shown for segment 2 ORFs of AejapNV1 and CxNV1 (GenBank MW226856.1). In **(A)** is shown the AejapNV1 segment 2 fORF, in **(B)** its rORF; in **(C)** the CxNV1 segment 2 fORF and in **(D)** its rORF. A1, B1, C1, and D1 show the pLDDT values of the five calculated models for each ORF. A2, B2, C2, and D2 show the five estimated output models superposed. Confidence ranking values for all models can be found in Suppl. Table 7. A3, B3, C3, and D3 show the model with the highest confidence value for each ORF with pLDDT values rendered in its tertiary structure. The pLDDT color scale is represented in **(E)**. All models were represented in the chart and in the structural superposition using the purple flare color palette, ranging from light purple (model 1) to dark purple (model 5). The pLDDT color scale can be read as: very low confidence (pLDDT < 50), low confidence (70 > pLDDT > 50), confident (90 > pLDDT > 70) and very high confidence (pLDDT > 90).

